# A modular live-cell biosensor for extracellular protease activity

**DOI:** 10.64898/2026.06.19.733436

**Authors:** Beatrice Ramm, Maia Weatherly, Jared Toettcher

## Abstract

Extracellular protease activity plays key roles in cell and tissue behavior, processing cell surface proteins, ligands and the extracellular matrix. Extracellular proteases can be subject to complex post-translational regulation, yet it remains challenging to quantify their activity in single cells over time. We present eNRGies (engineered neuregulin reporters as generalized indicators of extracellular shedding): modular, genetically encoded protease biosensors that translate extracellular cleavage into nuclear translocation of an intracellular (fluorescent) protein domain. We optimize the platform to report on protease activity in single cells on a timescale of minutes, and show it can be applied to soluble and cell-surface proteases including TEV protease, enterokinase, Factor Xa, MMP-9, and the sheddase ADAM17. We find ADAM17 activity can be transiently activated during mitosis and exhibit complex dynamics following EGF receptor stimulation. eNRGies biosensors enable observation of extracellular protease activity with high spatiotemporal resolution, and could be applied as synthetic biology scaffold to translate protease activity into customized cellular responses.

## Introduction

Extracellular proteases are key regulators of cell and tissue physiology. At the organismal scale, serine proteases initiate the blood clotting cascade and digest dietary protein. Within tissues, matrix metalloproteinases (MMPs) sculpt the cellular microenvironment by degrading extracellular matrix (ECM) components^1^. At the cellular scale, diverse cell signaling processes are regulated by proteases termed sheddases that trigger the proteolytic release of ligands or receptor ectodomains from the cell surface to activate cell signaling pathways^2^. A prominent class of sheddases are members of the A Disintegrin And Metalloprotease (ADAM) family of cell-surface proteases that release TNFα and growth factors from cells^3–6^ and initiate Notch-Delta signaling by cleaving Notch receptors^7^. Through these activities, extracellular proteases orchestrate biological processes including tissue remodeling and repair, organ morphogenesis, cell signaling, and wound healing. Extracellular proteases are also gaining prominence as potential targets for cancer therapeutics, due to their role in regulating the availability of growth factors and invasion through the extracellular matrix^2, 8^.

Due to their crucial physiological roles and the irreversibility of peptide bond cleavage, extracellular proteases are often tightly regulated. Their activity may be modulated by proteolytic processing, cofactor binding, oligomerization or post-translational modifications and in many cases is further controlled by regulatory and inhibitory proteins^9, 10^. Hence, protease abundance may not correlate with its activity. Yet measuring the activity states of extracellular proteases remains difficult. Traditionally, protease activity can be detected by zymography, the bulk conversion of labeled substrates delivered to a tissue.^11^ Similarly, end-point techniques have been used to analyze the amount of cleaved cell-surface substrates in the bulk cell media^4, 5, 12–15^. However, these techniques have limited spatiotemporal resolution because the fluorescent probes diffuse freely in the extracellular space as do the cleaved substrates which also accumulate slowly.

More recently, genetically encoded biosensors expressed on the cell-surface enabled the detection of extracellular cleavage events on a cellular level. Examples include a caged fluorogen activating protein^16^; a modified TNFα ligand whose cleavage generates a cryptic epitope that can be detected via fluorescently labeled nanobodies^17^; intensity or ratiometric biosensors based on substrates tagged with two fluorescent proteins such as neuregulin and TGFα^18–22^; and FRET-based biosensors for ADAM17 or MMPs^23–26^. However, while powerful, these techniques are typically non-modular as they are optimized for a single protease at a time. They also rely on a cleavage-induced change in fluorescence at the plasma membrane, which can be difficult to detect or assign to specific cells in crowded, tissues, and typically require normalization (e.g., comparing fluorescent signals before and after protease addition) to produce a quantifiable response. A high-precision biosensor that can be readily adapted to a wide range of proteases and allows for rapid detection and easy quantification in tissues is still lacking.

Here we present eNRGies, modular protease biosensors that can be readily adapted to multiple targets and report with a high dynamic range, a fast timescale of minutes, and fine single-cell spatial precision. eNRGies harness components of an endogenous protease-sensitive transmembrane protein – neuregulin – as the core of the biosensor, converting a cell-surface cleavage event into an intracellular response. We systematically optimize the biosensor’s extracellular domain, linkers, transmembrane sequence, and intracellular domain architecture, producing a biosensor architecture with a five-fold change in fluorescence localization to the nucleus upon protease cleavage. The biosensor’s quantitative response does not require normalization and can thus be used in both end-point assays and to measure protease dynamics over time. We show that eNRGies are modular and can be adapted to measure a variety of soluble and membrane-localized extracellular proteases. We apply eNRGies to measure ADAM17 activity in multiple cellular contexts, revealing transient pulses of protease activity during cell division in HEK293T cells and a protease-based positive feedback loop in epidermal growth factor receptor (EGFR) signaling in MCF10A cells. Our study points the way to a new generation of highly sensitive protease biosensors for drug discovery and characterization of dynamic, spatially complex extracellular states.

## Results

### Design and optimization of a biosensor for extracellular protease activity

Our overall goal was to develop a genetically encoded biosensor that relays an extracellular proteolytic event into an intracellular signal. Our design was inspired by regulated intramembrane proteolysis^27^ (RIP), a mechanism by which cells convert extracellular cleavage events into intracellular responses. In RIP, a primary cleavage event cuts the ectodomain in close proximity to the transmembrane domain and releases it into the extracellular space. In a second step, the remaining transmembrane domain undergoes cleavage by the ubiquitous intramembrane protease γ-secretase, releasing a short peptide into the extracellular space and the intracellular domain (ICD) into the cytosol. The protease γ-secretase is promiscuous with respect to substrate choice, with over 140 substrates identified to date^28^. However, the size of the transmembrane protein’s extracellular domain is thought to be a critical determinant of γ-secretase activity, with efficient cleavage requiring an ectodomain of less than 50 amino acids (AA)^29–31^, explaining why an initial ectodomain cleavage event is required. RIP is essential for processing of the amyloid precursor protein (APP)^32^, transmembrane growth factor precursors like neuregulin^33, 34^, and Notch receptor activation during Notch-Delta signaling^35^.

We hypothesized that the RIP mechanism could be harnessed as a biosensor for diverse extracellular proteases of interest (**Fig. 1a**). Our strategy involved replacing the primary cleavage site with a site for the protease of interest, appending fluorescent proteins to monitor cleavage via a change in fluorescence localization, and optimizing other features of the biosensor (e.g. linkers) to maximize the biosensor’s dynamic range. We used the epidermal growth factor (EGF) family member Neuregulin 1 type III (NRG1) (**Fig. 1b**) as a starting point, based on prior studies showing that its intracellular domain translocates to the nucleus following RIP and that a minimal construct lacking the NRG1 ectodomain is constitutively cleaved^33, 34^.

**Fig. 1:**
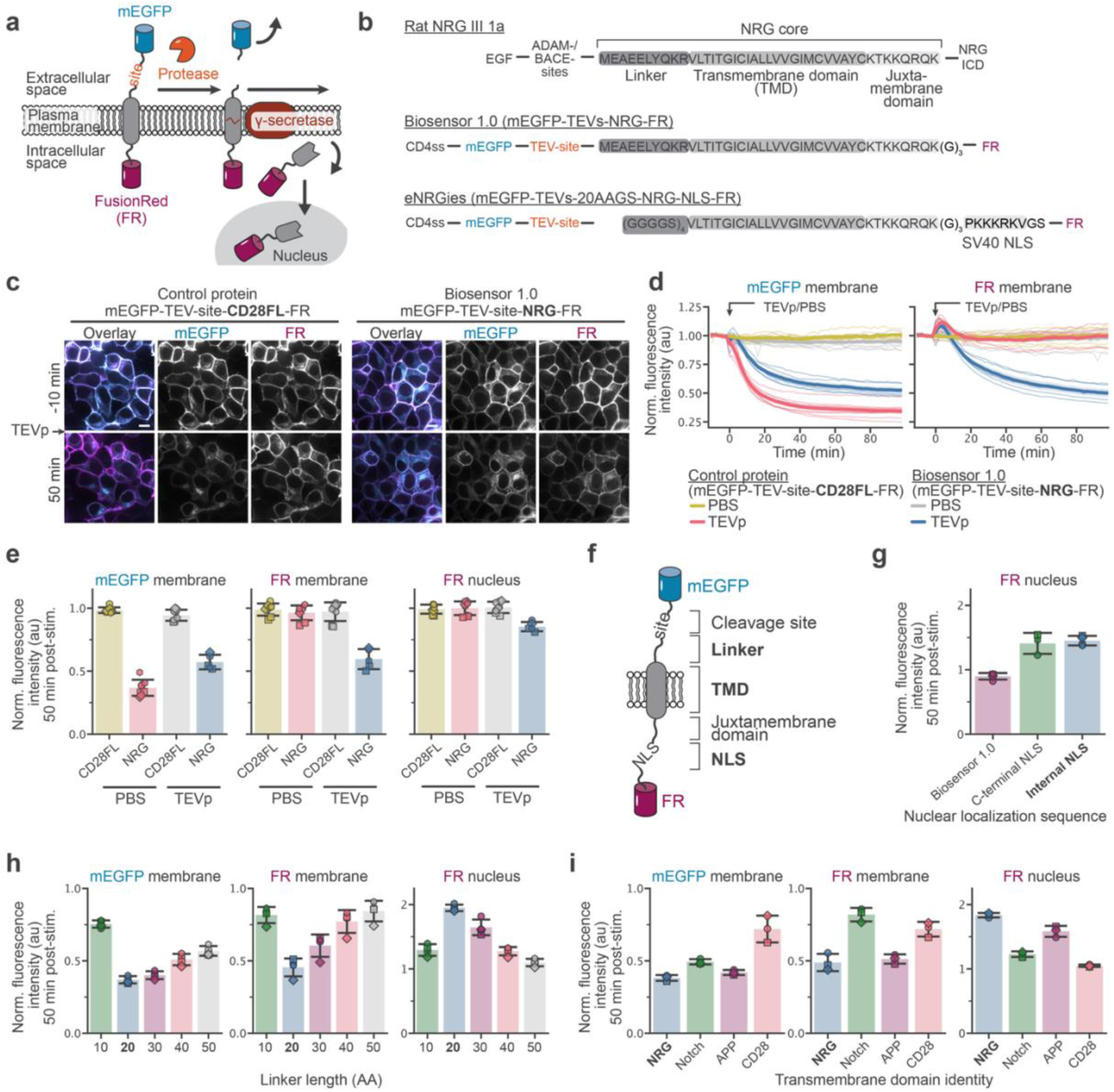
Design and optimization of a two-step biosensor for extracellular protease activity. **a**, Schematic of the biosensor mechanism. Protease activity liberates a bulky, extracellular domain (here mEGFP). The remaining transmembrane protein undergoes intramembrane proteolysis by γ-secretase releasing the ICD from the membrane, which then translocates into the nucleus. **b**, Schematic of the domain structure of the γ-secretase substrate Neuregulin 1 type III and of the initial (Biosensor 1.0) and final (eNRGies) biosensor designs. The extracellular domain is replaced by mEGFP, the endogeneous NRG cleavage sites (ADAM/BACE) are replaced by a TEVp cleavage site (TEV-site), and the intracellular domain is replaced by FR. **c**, Representative images of cells expressing the initial biosensor design or a control protein (full-length CD28 transmembrane protein) with the same extracellular and intracellular domains before or 50 min after exposure to 20 U TEVp. **d,** Time course of the normalized membrane mEGFP and FR fluorescence intensity in cells expressing the initial biosensor or the control protein exposed to PBS or 20 U TEVp. The bold lines and bands represent the mean ± SEM of traces from *N*(NRG_PBS_) = 8*, N*(CD28FL_PBS_) = 8, *N*(NRG_TEVp_) = 5, *N*(CD28FL_TEVp_) = 8 samples from 4 independent experiments, respectively. **e**, Normalized fluorescence intensity of mEGFP and FR on the membrane and FR in the nucleus quantified at 46-54 min post-stimulation with PBS or TEVp of cells expressing the initial biosensor or a full-length CD28 transmembrane protein. Bar and error bars represent the mean ± SD from *N*(NRG_TEVp_) = 5, *N*(CD28FL_TEVp_) = 8, *N*(NRG_PBS_) = 8, *N*(CD28FL_PBS_) = 8 independent experiments identified by marker, respectively. **f,** Schematic of the biosensor domains and features with those chosen for optimization highlighted in bold. **g-i**, Normalized fluorescence intensity of mEGFP and FR on the membrane and FR in the nucleus 46-54 min after TEVp exposure for (**g**) the initial design (Biosensor 1.0) and designs incorporating an additional NLS directly after the juxtamembrane domain or at the C-terminal end, (**h**) a biosensor with additional internal NLS in which the original NRG linker sequence has been replaced with a GS linker of varying length, and (**i**) a biosensor with additional internal NLS, 20-AA GS linker and different transmembrane domains. Bar and error bars represent the mean ± SD from *N*(Biosensor 1.0) = 4, *N*(C-terminal NLS) = 3, *N*(Internal NLS) = 4, *N*(10 AA) = 4, *N*(20 AA) = 4, *N*(30 AA) = 4, *N*(40 AA) = 4, *N*(50 AA) = 4, *N*(NRG) = 3, *N*(Notch) = 3, *N*(APP) = 3, *N*(CD28) = 3 independent experiments identified by marker, respectively. Scale bars, 10 μm.

Our initial biosensor design consisted of an mEGFP fluorescent extracellular domain (ECD), a protease cleavage site, the neuregulin core, and a FusionRed (FR) intracellular domain (ICD) (**Fig. 1b**). We chose the TEV protease (TEVp) cleavage site (TEV-site) as our initial target because of its high substrate specificity and ready availability^36^. The neuregulin core domain consists of a 10-amino acid linker sequence^37^, the NRG transmembrane domain containing the γ-secretase cleavage site, and an intracellular juxtamembrane domain. It is devoid of other annotated extracellular protease cleavage sequences. As a control, we used a similar construct harboring the full-length CD28 transmembrane protein (CD28FL) which is not a γ-secretase substrate. We generated monoclonal cell lines that also expressed a membrane-localized iRFP (myr-iRFP) and a nucleus-localized TagBFP (NLS-BFP) to easily segment the plasma membrane and nucleus using Cellpose^38^, enabling quantification of our biosensor’s membrane and nuclear fluorescence intensities across many cells (**Fig. S1**).

When we added a high dose of recombinant TEVp to cells expressing either our biosensor or the CD28 control construct, we observed rapid loss of the mEGFP membrane signal, indicative of its efficient cleavage and release into the medium (**Fig. 1c, d**). We also observed a decrease in membrane FR fluorescence over time for the biosensor, but not the CD28FL control construct, consistent with γ-secretase cleavage of the NRG core and release of the ICD from the membrane (**Fig. 1c, d, Movie S1**). In cells that were imaged concurrently but not exposed to TEVp, the fluorescence intensity of mEGFP and FR remained relatively constant, demonstrating that the loss of membrane mEGFP/FR signal was not due to photobleaching (**Fig. 1d**). Notably, we did not observe any accumulation of FR fluorescence in the nucleus, even though the juxtamembrane domain of NRG harbors a stretch of basic amino acids that are thought to function as an NLS^33^, suggesting that the cleaved ICD was rapidly degraded (**Fig. 1e**).

Nuclear biosensor accumulation is highly desirable because of the ease of quantification of nuclear signals, so we sought to optimize version 1.0 of our biosensor to improve its nuclear ICD accumulation. We targeted the ICD reporter, transmembrane domain sequence, and ectodomain linker as sites for potential improvement (**Fig. 1f**). First, to increase nuclear import of the ICD, we incorporated an additional NLS either near the juxtamembrane domain (internal NLS) or at the C-terminus (C-terminal NLS). Cells harboring the biosensor with these additional NLS variants showed a ∼50% increase in nuclear FR signal after incubation with TEVp (**Fig. 1g, Movie S2**). We proceeded with the internal NLS variant, as the C-terminal NLS variant exhibited more extensive ER localization, suggesting that it was trafficked less efficiently to the cell surface (**Fig. S2**).

We next hypothesized that varying the linker sequence between the TEV cleavage site and the NRG transmembrane domain could further optimize the efficiency of cleavage and biosensor response. We tested two types of linkers – a flexible GS linker and a helical EAAAK linker – with lengths ranging from 10 to 50 AAs (**Fig. 1h, Fig. S3, Movie S3**). With both linker types an interesting pattern emerged: FR nuclear intensity peaked for the 20-AA linkers compared to either shorter or longer linkers, with loss of membrane FR showing a similar trend **(Fig. 1h**). In contrast, membrane loss of extracellular mEGFP was pronounced for all linkers 20 AAs in length or longer (**Fig. 1h**). Our data is consistent with a model where linker length affects accessibility to both the TEVp and γ-secretase proteases. On one hand, a short 10-AAfor linker would restrict TEV protease from accessing its extracellular cleavage site, leading to low mEGFP cleavage. On the other hand, an excessively long linker would permit efficient TEVp cleavage but restrict access of the TEVp-cleaved substrate to γ-secretase, blocking FR redistribution from the membrane to the nucleus. A 20-AA long linker appears to balance these two effects producing an optimal biosensor response. We found that flexible linkers outperformed helical ones in all cases, and we proceeded with the 20-AA GS linker for all subsequent experiments (**Fig. S3**).

Finally, we tested whether alternative transmembrane domains (TMDs) might yield further biosensor improvements (**Fig. 1i, Movie S4**). We chose TMDs from the well-characterized γ-secretase substrates APP and Notch, as well as those of the non-substrate CD28. Each of the resulting constructs trafficked less efficiently to the cell surface than our original design harboring the NRG TMD (**Fig. S4**). The Notch construct, and to a lesser extent the APP construct, were not only localized on the membrane but also in the nucleus prior to simulation, and the CD28 construct showed extensive ER localization and only faint membrane localization. While the APP TMD was cleaved at a rate similar to the one from the NRG TMD (**Fig. S4**), we observed weak or nonexistent cleavage for the CD28 and Notch TMDs. Overall, we concluded that the superior surface localization and potent cleavage of the NRG-based core domain make this protein an excellent platform for our biosensor. We refer to our optimized biosensor architecture consisting of mEGFP, a protease cleavage site, a 20-AA GS linker, the NRG TMD, and the NLS-FR ICD, by the term “eNRGies”, for “<u>e</u>ngineered <u>n</u>euregulin reporters as <u>g</u>eneral indicators for <u>e</u>xtracellular <u>s</u>hedding” (**Fig. 1b**).

### Characterization of the biosensor

We next set out to quantitatively characterize the response of the eNRGies biosensor. We stimulated cells expressing TEVp-cleavable eNRGies with 0 to 20 U of recombinant TEVp and quantified membrane mEGFP, membrane FR and nuclear FR over time (**Fig. 2a, b, Movie S5**). Biosensor responses were qualitatively apparent from the mEGFP/FR overlay images, with a loss of membrane mEGFP fluorescence and near-complete redistribution of FR fluorescence from membrane to nucleus within 50 min in response to 20 U of TEVp (**Fig. 2a**). Quantifying membrane and nuclear fluorescence revealed that biosensor response kinetics and amplitudes varied as a function of TEVp dose (**Fig. 2a, b**).

**Fig. 2:**
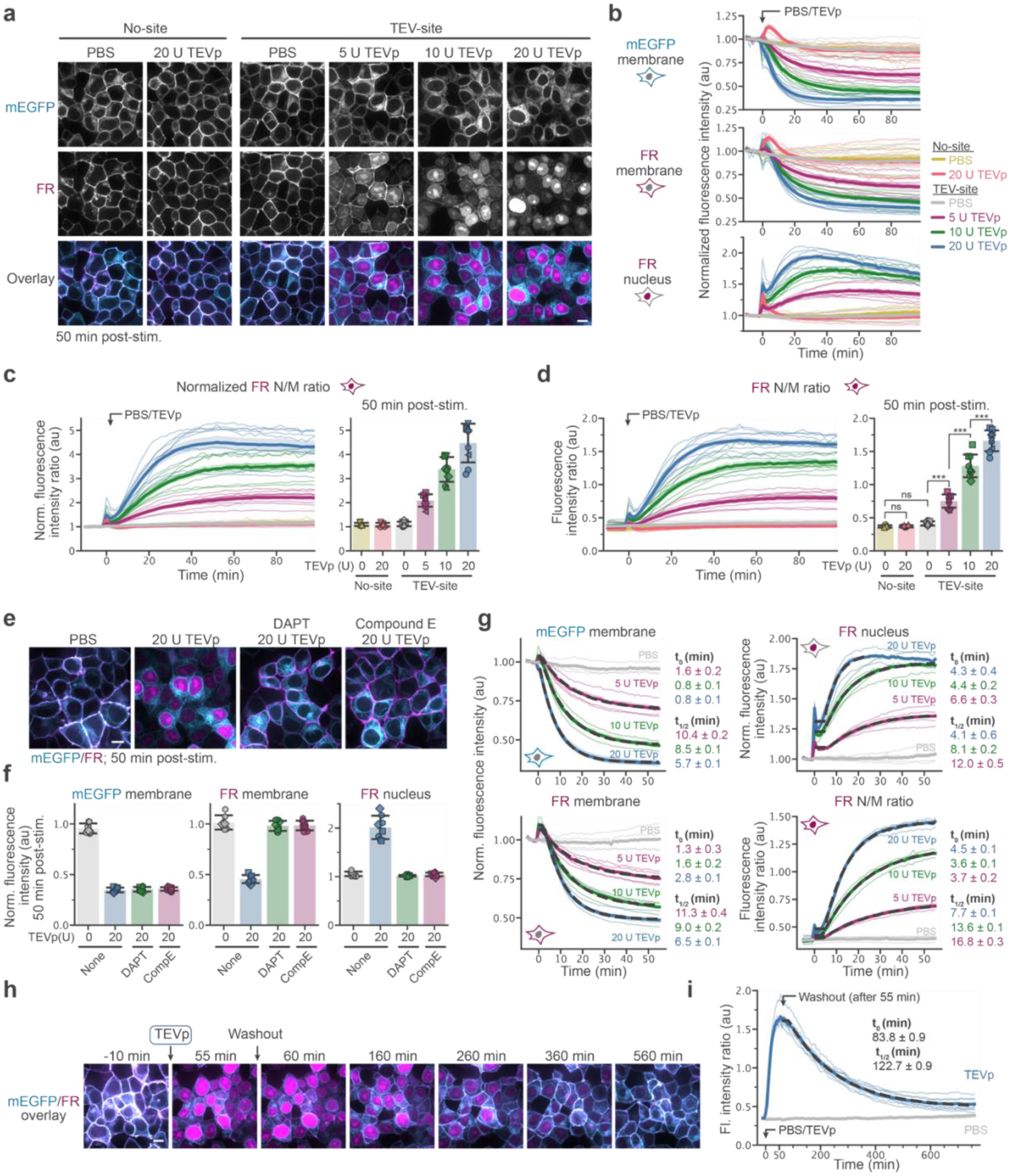
Characterization of the biosensor for extracellular protease activity. **a**, Representative images of cells expressing TEVp (TEV-site) or a site-less control eNRGies (No-site) 50 min after exposure to varying amounts of TEVp at time = 0. **b**, Normalized fluorescence intensity time courses of mEGFP and FR on the membrane and FR in the nucleus of cells expressing a cleavage site-less or TEVp eNRGies when exposed to varying amounts of recombinant TEVp at time = 0. **c, d** Quantification of FR N/M ratio either (**c**) normalized to the pre-stimulus value or (**d**) not normalized. Responses over time (left panels) and at a single time point 46-54 min post-TEVp exposure (right panels) are shown. TEVp doses are colored as indicated in **(b**). The bold lines and band represent the mean ± SEM of traces, bar and error bars represent the mean ± SD from *N*(No site - PBS) = 6, *N*(No site – 20 U TEVp) = 6, *N*(TEVs - PBS) = 8, *N*(TEVs – 5 U TEVp) = 8, *N*(TEVs – 10 U TEVp) = 9, *N*(TEVs – 520 U TEVp) = 9 independent samples from at least three independent experiments identified by marker, respectively. Statistical analysis was conducted using a two-sided Welch’s *t*-test ns > 0.05, \**P* ≤ 0.05, \*\**P* ≤ 0.01 and \*\*\**P* ≤ 0.001. **e**, **f**, Representative images (**e**) and normalized fluorescence intensity of mEGFP and FR on the membrane and FR in the nucleus (**f**) of cells expressing the TEVp eNRGies 46-54 min after exposure to 20 U TEVp that were pretreated with or without γ-secretase inhibitors DAPT and Compound E. Bar and error bars represent the mean ± SD from *N*(PBS) = 7, *N*(20 U TEVp) = 8, *N*(DAPT 20 U TEVp) = 8, *N*(Compound E 20 U TEVp) = 8 samples from 4 independent experiments identified by marker, respectively. **g**, Normalized fluorescence intensity of mEGFP and FR on the membrane and FR in the nucleus and the FR N/M ratio over time of a monoclonal cell line expressing TEVp eNRGies when exposed to varying amounts of recombinant TEVp at time = 0. Bold lines represent the mean of traces from *N*(PBS) = 4, *N*(5 U TEVp) = 4 *N*(10 U TEVp) = 4, *N*(20 U TEVp) = 3 independent experiments, respectively. Expontential fits to the response are shown in dark dashed lines with the obtained delay parameter t_0_ and half-life t_1/2_ indicated in the corresponding color. **h,** Representative images of cells expressing the TEVp biosensor that were exposed to 20 U TEVp at time = 0 and were washed after 55 min of incubation to remove TEVp. **i**, FR N/M ratio of cells expressing the TEVp biosensor that were exposed to PBS or 20 U TEVp at time = 0 and were washed after 55 min of incubation to remove PBS or TEVp. Bold lines represent the mean of traces from *N*(PBS) = 4, *N*(TEVp) = 8 independent experiments, respectively. Expontential fits to the response function are shown in dark dashed lines and the delay parameter and half-life are indicated. Scale bars, 10 µm.

We reasoned that the FR nuclear-to-membrane (N/M) fluorescence ratio could serve as an ideal single metric for quantifying biosensor responses. This metric is particularly useful because it only depends on a single fluorophore and because it quantifies a fluorescence ratio at two subcellular locations, which should be relatively independent of biosensor expression level and imaging conditions. When normalized to the cells’ initial state, this metric undergoes an approximately five-fold change between unstimulated and stimulated conditions (**Fig. 2c**). Importantly, the FR N/M fluorescence ratio does not require normalization to be interpretable and is highly reproducible between independent experiments, so its raw value can also be used in single-timepoint endpoint assays (**Fig. 2d**). Indeed, the N/M ratio increased with TEVp dose, and the N/M ratio in the absence of TEVp was comparable to the output of a biosensor that lacked a protease cleavage site (**Fig. 2a-d**).

While eNRGies biosensors are designed to report on extracellular protease activity, they also rely on secondary cleavage by γ-secretase and subsequent nuclear import of the FR fluorophore. Indeed, pretreatment with the γ-secretase inhibitors DAPT or Compound E prior to TEVp treatment confirmed that γ-secretase-dependent intramembrane proteolysis is essential for FR localization to the nucleus (**Fig. 2e, f, Movie S6**). Ideally, for eNRGies to reflect the real-time kinetics of the protease of interest, γ-secretase cleavage must be rapid relative to extracellular cleavage. To test whether this criterion is met, we titrated TEVp concentration and acquired time-series with one minute resolution in a monoclonal cell line harboring the TEVp eNRGies biosensor. We sought to compare the rate of mEGFP loss, which only depends on extracellular protease activity, with that of FR membrane loss, which also depends on γ-secretase. We found that these half-lives were within 1 min of one another across a wide range of TEVp doses, suggesting that γ-secretase operates highly efficiently. The increase of FR in the nucleus had a larger time-delay t_0_ than the membrane signal losses, but comparable half-lives. The FR N/M ratio is a ratio of two exponentials, and the resulting parameters are correlated but not identical to those of the FR fluorescence intensity signals (**Fig. 2g**). Overall, our data suggests that biosensor kinetics are primarily determined by extracellular protease activity and that biosensor redistribution is rapid, reporting on protease activity with an approximate timescale of ∼10 min.

Protease cleavage is irreversible. A protease biosensor’s post-stimulus return to baseline may thus be slow, as it depends on the degradation of the cleaved products and re-synthesis of the uncleaved biosensor. To determine the kinetics of reversibility, we exposed HEK293T cells harboring the TEVp-eNRGies biosensor to TEVp for 1 h, washed to remove the protease, and quantified the FR N/M ratio over time. We found that the N/M ratio returned to baseline with a half-life of 123 min, comparable to the half-life of the FR nuclear signal decay of 149 min (**Fig. 2h, Fig. S5, Movie S7**). While the half-life of the FR membrane signal could not be directly determined because of photobleaching during the multi-hour acquisition and slow maturation time of the protein, the recovery of mEGFP membrane fluorescence took much longer, with a half-life of about 252 min. This shows that recovery of the N/M ratio representing the biosensor output is dominated by the recovery of the nuclear signal. In principle, if faster kinetics are desired, expressing a destabilized variant of eNRGies could accelerate both the removal of cleaved components and the return to the uncleaved steady-state, at the expense of overall fluorescent brightness^39^.

### Extending the applicability of eNRGies

An ideal biosensor platform would exhibit a high degree of modularity, capable of responding to many different proteases, but also be compatible with a wide variety of extracellular and intracellular domains to enable different applications. We thus tested whether eNRGies biosensors are modular with respect to extracellular domains and intracellular reporters. We first replaced the extracellular mEGFP domain with an alternative non-fluorescent domain, freeing up an additional fluorescent channel (**Fig. 3a**). We chose the reporter protein SEAP, as it is known to traffic well, and its enzymatic activity in the extracellular medium can be used to monitor overall shedding by quantifying the enzymatic turnover of a chromogenic substrate^5, 12, 13^. The biosensor showed similar maximum responses when we replaced mEGFP with SEAP, albeit with slightly slower kinetics (**Fig. 3b, c, Fig. S6**). This may be explained by the more restricted access of TEVp to the cleavage site, as SEAP is a bulkier protein than mEGFP (55 vs 27 kDa).

**Fig. 3:**
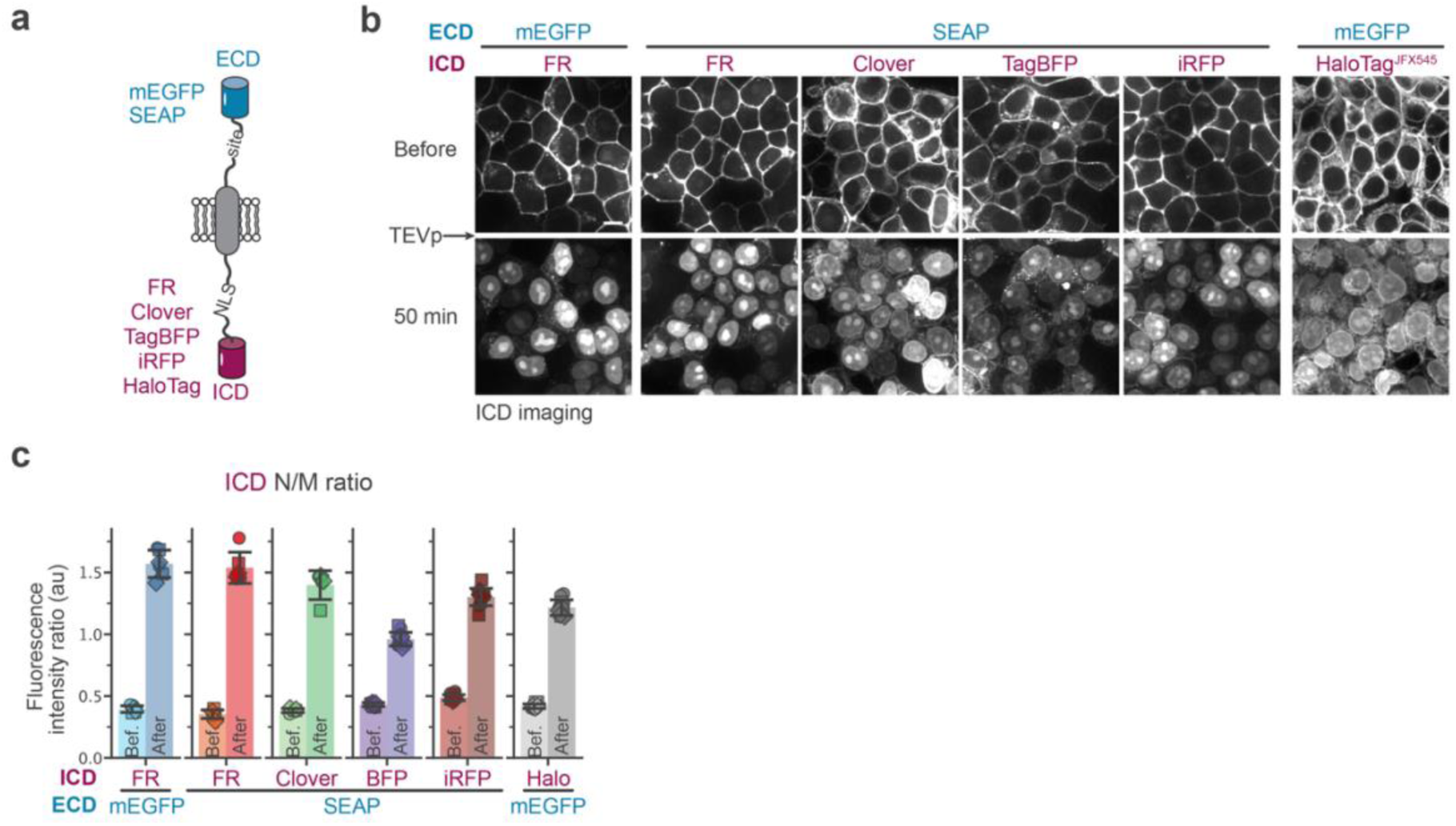
eNRGies are a modular biosensor platform across multiple ECDs and ICDs. **a,** Schematic of eNRGies highlighting the interchangeable bulky ECD and the ICD. **b**, Representative images of cells acquired in the ICD fluorescence channel expressing the TEVp eNRGies with different ECD and ICD domains before or 50 min after exposure to 20 U of TEVp at time = 0. eNRGies with a HaloTag ICD were labeled with a JFX554 HaloTag ligand before image acquisition. **c**, N/M ratio of the ICD fluorescence reporter of cells expressing a TEVp eNRGies before and 50 min after exposure to 20 U recombinant TEVp at time = 0. Bar and error bars represent the mean ± SD from *N*(mEGFP-FR) = 6, *N*(SEAP-FR) = 6, *N*(SEAP-Clover) = 5, *N*(SEAP-BFP) = 12, *N*(SEAP-iRFP) = 12, *N*(mEGFP-HaloTag) = 12 independent samples and three independent experiments identified by marker, respectively. Scale bars, 10 µm.

Since our N/M ratio readout is a ratio of subcellular localization intensities for a single fluorophore, it should not depend on the specific identity of the fluorescent reporter. We tested several fluorescent reporters in the context of the ICD: TagBFP, Clover, iRFP, and the HaloTag fluorescent indicator that can be used with a variety of chemical Halo dyes (**Fig. 3a-c, Fig. S6, Movie S8**). The HaloTag variant was expressed in the context of the mEGFP extracellular domain; all others were cloned with a SEAP extracellular domain. We swapped the fluorescent proteins used as constitutive nuclear and membrane markers for segmentation when needed. All reporters showed clear translocation from the membrane to the nucleus after TEVp addition (**Fig. 3b, c**). The N/M ratio was similar when FR or Clover were used as reporter, with slightly lower N/M ratios observed for HaloTag, TagBFP and iRFP, potentially due to reporter-specific differences in ICD nuclear import or degradation. Overall, these results demonstrate that eNRGies are versatile reporters that exhibit similar responses in the context of various intracellular and extracellular domains.

### eNRGies are versatile biosensors for soluble extracellular proteases

We next tested whether eNRGies biosensors are modular with respect to multiple recombinant proteases (**Fig. 4a**). We designed biosensors for two additional widely used recombinant proteases: Factor Xa (FXa) and Enterokinase (EK). This was done by replacing the TEVp cleavage site with cleavage sites for these other proteases. We transfected cells with one of the three biosensors and incubated with high concentrations of the cognate protease, observing robust cleavage in all three cases (**Fig. 4b, c, Movie S9**). All three biosensors showed a baseline N/M ratio of 0.4 and a maximum activation of ∼1.7 for their cognate protease, suggesting that the quantitative response of eNRGies is comparable between different proteases.

**Fig. 4:**
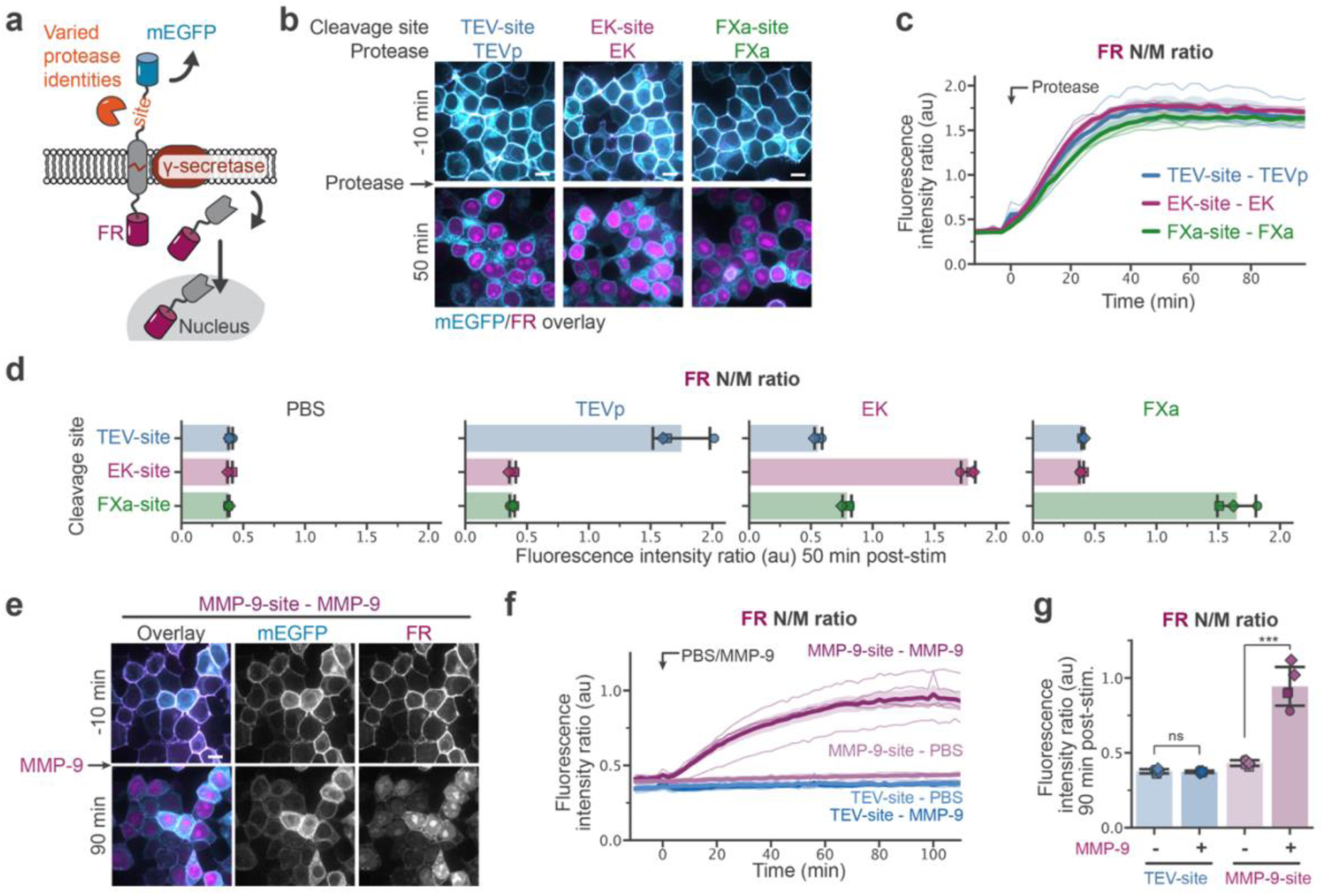
eNRGies can report on the activity of various soluble extracellular proteases. **a,** Schematic of the eNRGies biosensor highlighting the cleavage site that can be swapped out to detect various soluble proteases. **b**, Representative images of cells expressing eNRGies biosensors with a TEV-site, EK-site and FXa-site before and after exposure to 20 U TEVp, 32 U EK and 1 µg FXa at time = 0, respectively. **c**, FR N/M ratio time course of cells expressing eNRGies biosensor with a TEV-site, EK-site and FXa-site exposed to 20 U TEVp, 32 U EK and 1 µg FXa, respectively, at time = 0. **c,** FR N/M ratio over time for cells treated as in **b**. Bold lines and band represent the mean ± SEM of traces from three independent experiments. **d,** FR N/M ratio at 50 min post-stimulation for cells expressing eNRGies biosensors with TEV-site, EK-site, and FXa-site and treated with proteases in all combinations as in **c.** Bar and error bars represent the mean ± SD from three independent experiments identified by marker. **e,** Representative images of cells expressing an eNRGies biosensor with MMP-9-site before and after exposure to PBS or 28 U MMP-9 at time = 0. **f, g**, FR N/M ratio over time (**f**) and at 90 min post-stimulation (**g**) for cells expressing eNRGies biosensors with a TEV-site or MMP-9-site and exposed to PBS or MMP-9 at time = 0 as in **e**. The bold lines and band represent the mean ± SEM of traces, bar and error bars represent the mean ± SD from 5 samples from three independent experiments identified by marker. Statistical analysis was conducted using a two-sided Welch’s *t*-test. ns > 0.05, \**P* ≤ 0.05, \*\**P* ≤ 0.01 and \*\*\**P* ≤ 0.001. Scale bars, 10 µm.

To assess specificity, we also measured responses for all pairwise combinations of biosensors and proteases (**Fig. 4d**). While TEVp and FXa were only able to activate their cognate biosensor, EK also drove weak activation of the FXa biosensor. This result is consistent with prior reports that EK exhibits low site specificity, cleaving a broad range of sequences especially if they are accessible^40^. These results highlight that the biosensor can be readily reprogrammed to other proteases and suggest that eNRGies could be used to map the specificity of a protease of interest by constructing biosensors harboring cleavage sites variants and measuring quantitative responses.

As proof of concept that eNRGies can also respond to more physiologically relevant extracellular proteases, we also constructed an eNRGies biosensor for the extracellular matrix metalloproteinase MMP-9 that incorporates a consensus substrate peptide for MMP-9 cleavage^24,41^. Addition of recombinant MMP-9 to cells resulted in activation of the MMP-9 biosensor, whereas the TEVp biosensor was not cleaved (**Fig. 4 e-g, Movie S9**). We observed slower kinetics of biosensor accumulation and a lower maximum response amplitude for MMP-9 compared to other proteases (**Fig. 4c, f**), suggesting that cleavage was less efficient than for the other recombinant proteases tested. Taken together, these data suggest that eNRGies can be rapidly developed for a wide range of soluble extracellular proteases.

### Adapting eNRGies to the ADAM17 cell-surface protease

With our eNRGies biosensors in hand, we set out to investigate the activity dynamics of the ADAM17 protease in human cells. ADAM17 and a related homologue ADAM10 are among the best-studied members of the A Disintegrin And Metalloprotease (ADAM) family, a family of cell surface-expressed proteases that regulate the activity of a variety of cell signaling pathways^3–5, 7, 42,43^. For example, Notch/Delta signaling is initiated upon proteolysis of the Notch receptor mainly by the ADAM10 cell-surface protease^7^, and ADAM17 is responsible for processing many membrane-tethered pro-ligands into their soluble, active forms including tumor necrosis factor α (TNFα)^6^ and multiple members of the EGF (epidermal growth factor) family of epidermal growth factor receptor (EGFR) ligands^3–5^.

We first set out to construct an eNRGies biosensor harboring a peptide sequence that is specifically cleaved by ADAM17, a challenging goal due to the overlapping substrate specificity between ADAM10 and ADAM17. A recent *in vitro* screen^44^ identified a potential set of specific substrates: “TNFtide” and “TACEtide”, which are expected to be ADAM17-specific; “TENtide”, which is expected to be ADAM10 specific; and “ADAMtide”, which is expected to be cleaved mainly by ADAM10 and to a lesser extent by ADAM17. We generated eNRGies biosensors incorporating each cleavage site and introduced them into a panel of HEK293T cell lines: a parental line (WT) and derivatives where ADAM10 (10 KO), ADAM17 (17KO), or both (DKO) were knocked out^17^. Note that the parental HEK293T cell line used to generate these knockout lines is distinct from the standard HEK293T line used in our other experiments.

ADAM10 exhibits constitutive activity that may be further stimulated by mechanical forces/ionomycin^45, 46^, whereas ADAM17 can be induced by stimuli such as growth factors, phorbol ester 12-myristate 13-acetate (PMA), and anisomycin^4, 5, 47, 48^ (**Fig. 5a**). We thus examined the response of our candidate biosensors in the presence or absence of PMA (**Fig. S7**). All sensors seemed to only respond to ADAM10 and ADAM17 as their baseline and PMA-dependent response in DKO cells was comparable to the control sensor. The ADAMtide sensor exhibited PMA-dependent cleavage in 10KO cells and elevated PMA-independent cleavage in 17KO cells, consistent with its role as a substrate for both ADAM10 and ADAM17. In contrast, the sensors with the putative ADAM17 substrates TNFtide and TACEtide still exhibited elevated cleavage in 17KO cells compared to the control sensors, suggesting that they are not solely cleaved upon PMA simulation (**Fig. S7**). The ADAM10-specific TENtide exhibited similar qualitative trends to ADAMtide, but was not significantly cleaved upon PMA stimulation. All four peptides produced high biosensor activity in WT cells that could not be further stimulated by PMA addition. We therefore concluded that each peptide is likely to cross-react with multiple cell-surface proteases on cells, and that none are suitable as ADAM17-specific protease biosensors.

**Fig. 5:**
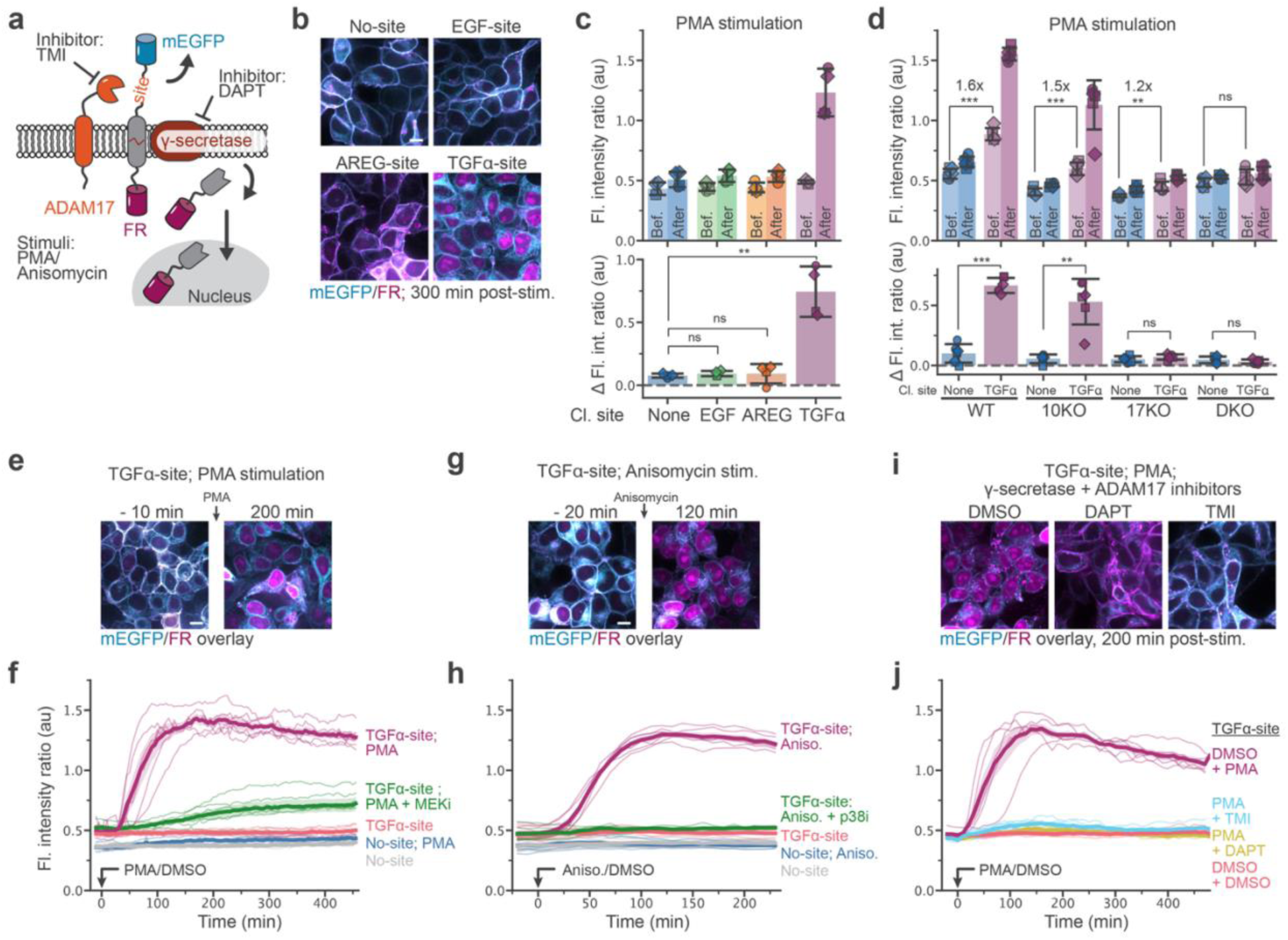
eNRGies reports on the activity of the sheddase ADAM17. **a,** Schematic depicting ADAM17 and eNRGies with extracellular mEGFP and intracellular FR, highlighting stimulatory and inhibitory agents of ADAM17 and γ-secretase. **b, c** Representative fluorescence images (**b**) and FR N/M ratio (top) and change in FR N/M ratio (bottom) (**c**) of cells expressing eNRGies with no cleavage site (No-site) or the proximal cleavage site of EGF, AREG or TGFα 5 min before and 290-310 min after stimulation with PMA. Bar and error bars represent the mean ± SD from 4 samples from three independent experiments identified by marker. **d**, FR N/M ratio (top) and change in FR N/M ratio (bottom) of WT HEK293T (WT), ADAM10KO (10KO), ADAM17KO (17KO) and double ADAM10/ADAM17KO (DKO) cells expressing eNRGies with no site or the TGFα proximal cleavage site before (-15/-7.5 min) and after stimulation (202.5-292.5 min) with PMA. Bar and error bars represent the mean ± SD from *N*(WT_No-site_) = 6, *N*(10KO_No-site_) = 5, *N*(17KO_No-site_) = 6, *N*(DKO_No-site_) = 6, *N*(WT_TGFα-site_) = 6, *N*(10KO_TGFα-site_) = 6, *N*(17KO_TGFα-site_) = 6, *N*(DKO_TGFα-site_) = 6 samples and three independent experiments identified by marker. (**c,d**) Statistical analysis was conducted using a two-sided Welch’s *t*-test. ns > 0.05, \**P* ≤ 0.05, \*\**P* ≤ 0.01 and \*\*\**P* ≤ 0.001. Fold change of the mean values is indicated where appropriate. **e,** Representative fluorescence images of cells expressing TGFα-site eNRGies before and after stimulation with PMA. **f**, FR N/M ratio time course of cells expressing site-less or TGFα-site eNRGies that were stimulated with DMSO or PMA at t = 0 and TGFα-site eNRGies expressing cells pretreated with MEK inhibitor (MEKi) and stimulated with PMA at t = 0. **g**, Representative fluorescence images of cells expressing TGFα-site eNRGies before and after stimulation with anisomycin. **h**, FR N/M ratio time course of cells expressing site-less or TGFα-site eNRGies that were stimulated with DMSO or anisomycin at t = 0 and TGFα-site eNRGies expressing cells pretreated with p38 inhibitor (p38i) and stimulated with anisomycin at t = 0. **i**, Representative fluorescence images of cells expressing TGFα-site eNRGies that were pretreated with DMSO, the γ-secretase inhibitor DAPT or the ADAM17 inhibitor TMI after stimulation with PMA. **j**, FR N/M ratio time course of cells expressing TGFα-site eNRGies that were pretreated with DMSO, DAPT or TMI and stimulated with DMSO or PMA at t = 0. **f, h, j** Bold lines and band represent the mean ± SEM of traces from 6 samples from three independent experiments. Scale bars, 10 µm.

We next tested peptides from natural proteins that are thought to be ADAM10 or ADAM17 substrates: the proximal cleavage sites of EGFR ligand transmembrane precursors transforming growth factor alpha (TGFα), amphiregulin (AREG) and epidermal growth factor (EGF)^5^ (**Fig. 5b, c, Fig. S8**). We incorporated these peptides with a shorter 10-AA linker into eNRGies, since they occur only 5-10 AAs away from the transmembrane domain in their native pro-ligand contexts, and because we reasoned that shorter linkers would not impair γ-secretase activity. While the sensors with the EGF and AREG-derived sequences failed to respond to PMA stimulation, the sensor with the TGFα-derived sequence exhibited both low baseline activity and potent PMA-induced response in standard HEK293T cells (**Fig. 5b, c, Fig. S8a**). We were surprised that our AREG-based candidate biosensor did not respond to PMA despite AREG’s status as a classic ADAM17 substrate, a discrepancy that may be explained by mis-annotation of the cleavage site^49^.

We proceeded to test the TGFα-derived eNRGies in the ADAM knockout cell lines (**Fig. 5d**, **Fig. S8b**). The TGFα-derived eNRGies basal activity in DKO cells was indistinguishable from that of the control sensor, and showed increasing basal activity compared to the control in 10KO, 17KO and WT cells. Hence, the biosensor captures ADAM17 basal activity but also seems to be cleaved at a very low level by ADAM10. The TGFα-based eNRGies showed a potent change from baseline for both WT and 10KO cells upon PMA stimulation, but no significant change upon PMA stimulation in 17KO and DKO cells (**Fig. 5d**, **Fig. S8b**). We noted that the level of pre-stimulus cleavage was substantially higher in the parental HEK293T cell line compared to our standard HEK293Ts and all knockout lines, suggesting that this cell line exhibits a high basal level of ADAM protease activity. Based on its potent PMA-stimulated response in all ADAM17-expressing cell lines tested and low baseline response in 17KO cells, we selected the TGFα-derived peptide for further characterization as a potential ADAM17 eNRGies biosensor.

To further characterize the TGFα-based eNRGies biosensor, we measured its response to a wide variety of ADAM17 activating and inhibiting stimuli. We found that two ADAM17-activating stimuli, PMA and anisomycin, drove robust biosensor cleavage within ∼1 h after stimulation, whereas DMSO-treated cells and PMA-or anisomycin treated cells that expressed an uncleavable biosensor all failed to respond (**Fig. 5e-h**, **Fig. S9a-d, Movie S10, S11**). Prior studies have shown that ADAM17 activation depends on MEK and p38 signaling in the case of PMA and anisomycin, respectively^48^. Indeed, we found that the MEK inhibitor PD0325901 and p38 inhibitor SB203580 blocked cells from mounting a biosensor response (**Fig. 5f, h**). Moreover, treatment with the ADAM17 inhibitor TMI-1 (TMI)^50, 51^ and the γ-secretase inhibitor DAPT was sufficient to abolish the PMA-induced response (**Fig. 5i, j, Fig. S9e-f, Movie S12**). Based on these results, we determined that the eNRGies construct incorporating the TGFα cleavage site is a *bona fide* ADAM17 eNRGies biosensor.

### eNRGies reveals spontaneous ADAM17 dynamics in HEK293T

To obtain a first high-resolution portrait of ADAM17 activity at single-cell resolution, we generated clonal HEK293T cell lines harboring the ADAM17-eNRGies biosensor or a control biosensor lacking the TGFα cleavage site; we also generated a clonal ADAM17-eNRGies biosensor line in the 17KO cell line. Moreover, we modified our image analysis pipeline to compute the FR N/M ratio of individual cells in the field of view over time. Imaging and quantification of the 293T ADAM17 eNRGies cell line revealed striking heterogeneity in the biosensor activity with a skewed distribution indicative of higher ADAM17 activity in a subset of cells (**Fig. 6a**). This heterogeneity was not observed in ADAM17KO cells or cells expressing a control biosensor lacking the TGFα site (**Fig. 6a**). This data suggests that ADAM17 activity is heterogenous across a population of genetically identical cells, with a subset of cells exhibiting higher rates of ADAM17 protease activity than the baseline activity leading to biosensor cleavage.

**Fig. 6:**
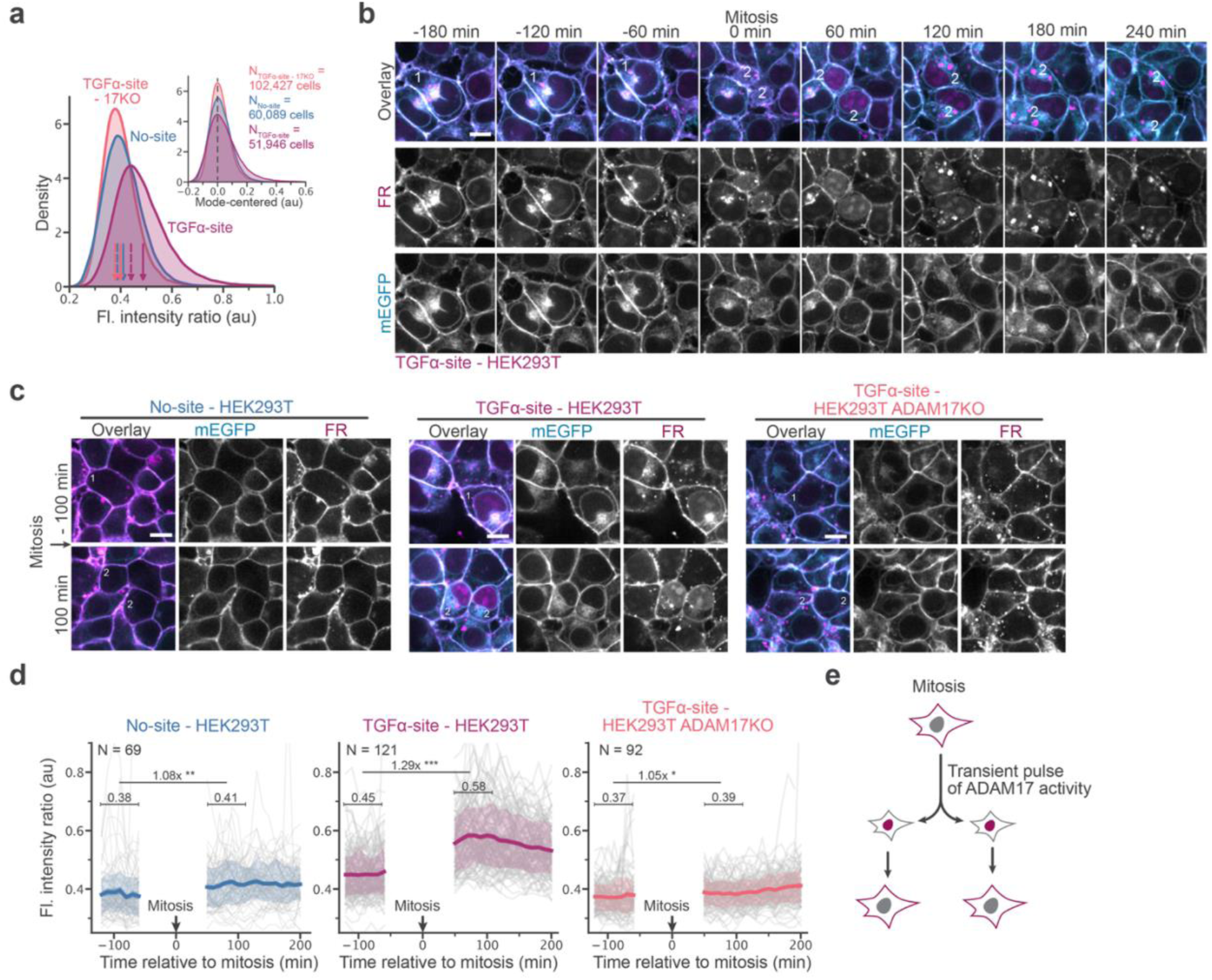
ADAM17 eNRGies reveals mitotic ADAM17 activity dynamics in HEK293T cells. **a,** Probability density distributions of the FR N/M ratio of individual HEK293T cells expressing a noncleavable eNRGies (No-site) or TGFα-site eNRGies and of ADAM17KO HEK293T cells expressing the TGFα-site eNRGies imaged over 600 min at 10 min/frame. Solid and dahsed colored arrows indicate the mean and mode. Inset: Mode-centered distribution of the FR N/M ratio. *N*(TGFα-site -17KO) = 18 FOV, 102,427 cells; *N*(No-site) = 9 FOV, 60,089 cells; *N*(TGFα-site) = 9 FOV, 51,946 cells from three independent experiments. **b**, Representative time-series of a HEK293T cell expressing TGFα-site eNRGies undergoing mitosis. Time 0 is defined as the time when mitosis is accomplished and the first occurrence of two daughter cells in the FOV. **c**, Representative fluorescence images of HEK293T cells expressing a noncleavable eNRGies or TGFα-site eNRGies and of ADAM17KO HEK293T expressing the TGFα-site eNRGies 100 min before and 100 min after cell division is finalized and two daughter cells are formed. **d,** Average FR N/M ratio of individual cells tracked from 120 min before to 200 min after mitosis of the cell types in **c**. Thin gray lines are traces of *N* individual cells, bold colored lines, dark/light colored bands represent the mean ± SEM and the 16th–84th percentile range, respectively, from 6 samples from three independent experiments. The 6 frames before and 5 frames after division were not plotted to exclude segmentation artifacts. Pre- and post-division mean values denoted over brackets were calculated over 6 frames with the fold change indicated. Statistical significances were determined by a paired two-sided Student’s *t*-test: ns > 0.05, \**P* ≤ 0.05, \*\**P* ≤ 0.01 and \*\*\**P* ≤ 0.001. **e**, Schematic of how transient ADAM17 acitivity during mitosis leads to biosensor activity. Scale bars, 10 µm.

How does ADAM17 activity vary over time in HEK293T cells? To address this question, we performed time-lapse imaging to monitor the accumulation and loss of eNRGies biosensor activity in single cells over time. Strikingly, we found that ADAM17 eNRGies biosensor activity was tightly correlated with cell cycle progression, with a transient peak of biosensor activity just following mitosis (**Fig. 6b**). Prior to mitosis, nuclear FR remained low, indicative of a low rate of ADAM17 cleavage. Following mitosis and upon nuclear envelope reformation, we observed increased nuclear FR intensity that gradually decreased over subsequent hours. Quantification of biosensor activity during individual mitotic events in all three cell lines revealed that the mitotic increase in biosensor activity was not present in ADAM17 knockout cells or cells expressing a site-less control biosensor, suggesting that it represented a *bona fide* pulse of ADAM17 activity associated with mitosis (**Fig. 6c-e, Movie S13**). Taken together, these data suggest that ADAM17 activity is transiently upregulated during mitosis in HEK293T cells (**Fig. 6e**). More broadly, our data demonstrate that cell-surface protease activity can be regulated with high spatiotemporal precision by endogenous cellular processes on a ∼1 h timescale.

### eNRGies reveals ADAM17 dynamics in EGF stimulated MCF10A monolayers

We next sought to apply our ADAM17 eNRGies biosensor in a cellular context where ADAM17 is thought to participate in complex spatiotemporal signaling. We turned to MCF10A human breast epithelial cells, where prior studies have found that ADAM17 is required for generating local tissue-scale EGFR/ERK signaling responses to BRAF oncogene activation and apoptotic events^52, 53^. Evidence also suggests that the EGFR/ERK pathway also feeds back to trigger ADAM17 cleavage: EGFR activation has been shown to increase shedding by ADAM17 in multiple epithelial cell contexts^20, 21, 23, 54^ and lung cancer cells harboring KRAS mutants exhibit increased shedding of IL-6R and EGFR ligands^55, 56^. This leads to the possibility of a feedback loop where EGFR stimulation increases ADAM17 activity, which in turn drives increased shedding of EGFR ligands (**Fig. 7a**), though the single-cell dynamics of these regulatory connections have not been measured.

**Fig. 7:**
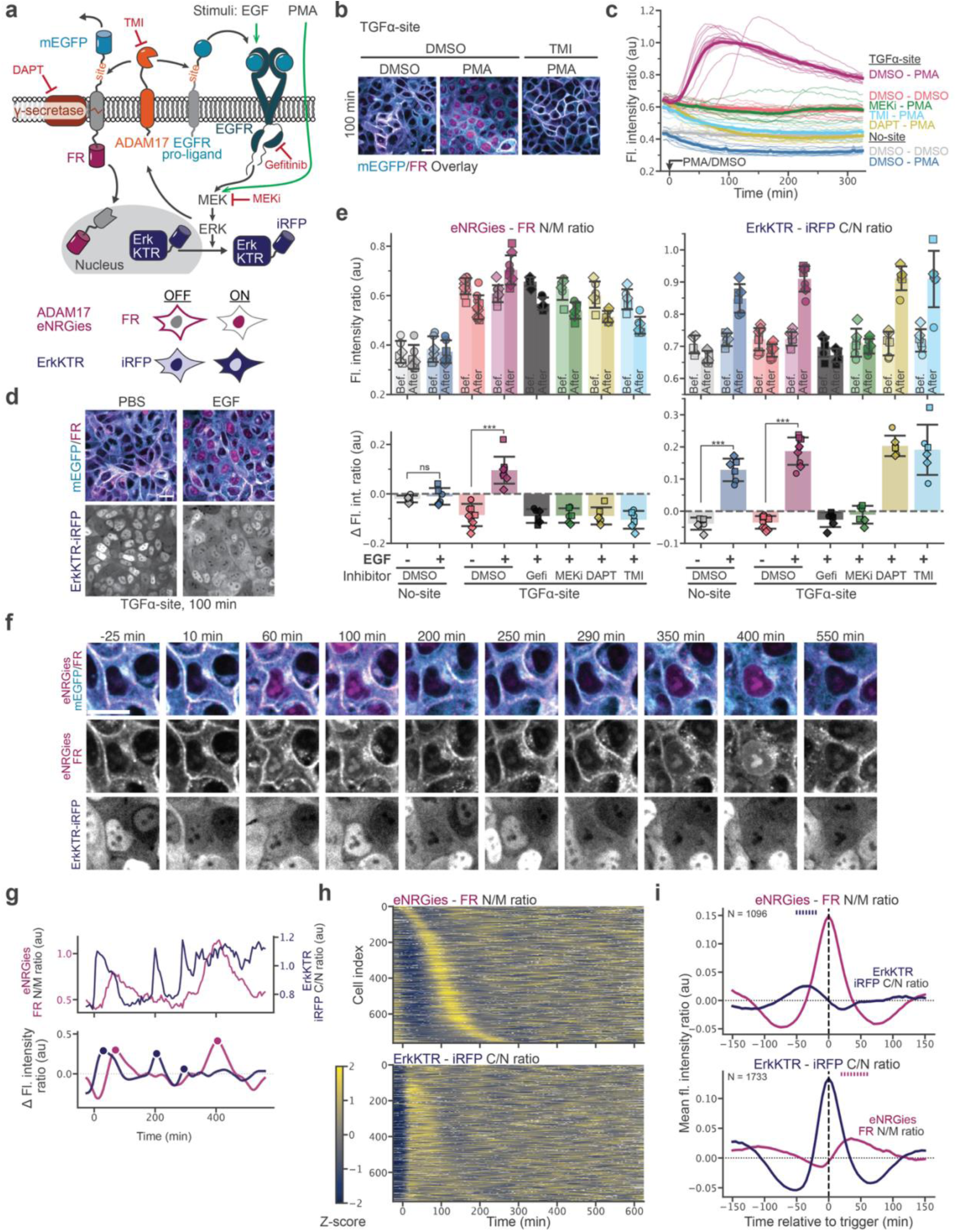
eNRGies biosensor reveals dynamics of ADAM17 activity in response to EGF stimulation. **a,** Schematic depicting ADAM17, EGFR pro-ligands, EGFR and their inhibitory and stimulating agents and the biosensors ADAM17 eNRGies and ErkKTR-iRFP highlighting their localization when ADAM17 and ERK activity is high (“ON”) or low (“OFF”). **b,** Representative fluorescence images of MCF10A cells expressing ADAM17 eNRGies pretreated with DMSO or TMI 100 min after stimulation with DMSO or PMA. **c,** FR N/M ratio time course of MCF10A cells expressing siteless or TGFα-site eNRGies that were pretreatd with DMSO, MEK inhibitor (MEKi), DAPT or TMI and stimulated with DMSO or PMA at t = 0. Bold lines and band represent the mean ± SEM of traces from *N*(No-site_DMSO_) = 8, *N*(No-site_PMA_) = 8, *N*(TGFα-site_DMSO_) = 12, *N*(TGFα-site_PMA_) = 12, *N*(TGFα-site_MEKi-PMA_) = 8, *N*(TGFα-site_DAPT-PMA_) = 8, *N*(TGFα-site_TMI-PMA_) = 8 samples and four independent experiments. **d**, Representative fluorescence images of MCF10A cells expressing ADAM17 eNRGies pretreated with DMSO 100 min after stimulation with PBS or EGF. **e**, eNRGies FR N/M ratio and ErkKTR-iRFP C/N ratio (top) and change in FR N/M ratio and iRFP C/N ratio (bottom) of MCF10A cells expressing eNRGies with no site or the TGFα-site and the ErkKTR-iRFP before (-25 to -5 min) and after stimulation (100-200 min) with PBS or EGF at t = 0 that were pretreated with DMSO, Gefitinib (Gefi), MEK inhibitor (MEKi), DAPT or TMI. Bar and error bars represent the mean ± SD from *N*(No-site_PBS_) = 6, *N*(No-site_EGF_) = 6, *N*(TGFα-site_PBS_) = 10, *N*(TGFα-site_EGF_) = 9, *N*(TGFα-site_Gefi-EGF_) = 8, *N*(TGFα-site_MEKi-EGF_) = 6, *N*(TGFα-site_DAPT-EGF_) = 6, *N*(TGFα-site_TMI-EGF_) = 6 samples and three independent experiments identified by marker. Statistical analysis was conducted using a two-sided Welch’s *t*-test. ns > 0.05, \**P* ≤ 0.05, \*\**P* ≤ 0.01 and \*\*\**P* ≤ 0.001. **f**, **g**, Representative fluorescence images (brightness and contrast not comparable across time points) (**f**) and single-cell fluorescence intensity ratio traces (**g**) of an MCF10A cell expressing ADAM17 eNRGies and ErkKTR-iRFP showing activity pulses when exposed to EGF at t = 0. Top panel shows raw flourescence intensity traces, lower panel shows smoothed and detrended traces with detected peaks marked (filled circles). **h**, Z-scored eNRGies FR N/M ratio and ErkKTR-iRFP C/N ratio per cell sorted according to the first peak in the eNRGies channel for 763 cells. White color indicate gaps in the tracking. **i**, Peak-triggered average of eNRGies FR N/M ratio (top) and ErkKTR-iRFP C/N ratio (bottom) peaks showing the aligned response of the non-triggered sensor. Shading indicates SEM, and N is the number of peak events averaged. Ticks indicate time points where the peak-triggered average of the co-recorded signal exceeds 70% of its peak amplitude. Scale bars, 20 µm.

To validate that our ADAM17 eNRGies biosensor was functional and specific for ADAM17 protease activity in MCF10A cells, we generated monoclonal cell lines harboring the ADAM17 eNRGies biosensor as well as nuclear and membrane markers for segmentation and quantification; an analogous clonal cell line expressing a non-cleavable eNRGies scaffold was generated as a negative control. MCF10As expressing the ADAM17 eNRGies biosensor responded to PMA treatment with redistribution of FR fluorescence from membrane to nucleus, and PMA-induced biosensor cleavage was also prevented by pharmacological inhibition of ADAM17, MEK, or γ-secretase (**Fig. 7a-c**, **Fig. S10a, b, Movie S14**). Cells harboring the site-less biosensor or treated with protease inhibitors exhibited a gradual decrease in FR N/M ratio after PMA treatment, possibly due to increased synthesis of the biosensor and membrane localization upon PMA stimulation (**Fig. 7c, Fig. S10a, b**). Overall, our results were strongly reminiscent of what was observed in HEK293T cells, with a similar ∼100 min timescale of PMA-stimulated biosensor activity (**Fig. 5e**, **Fig. 7c**), demonstrating that our ADAM17 eNRGies biosensor responds with comparable kinetics across multiple cellular contexts.

We then set out to investigate the interplay between ADAM17 and ERK activity dynamics after stimulation with the EGFR ligand EGF. We generated additional clonal cell lines harboring either the ADAM17 or site-less eNRGies biosensors, an NLS-BFP nuclear marker, and an iRFP-tagged biosensor of ERK activity (ErkKTR-iRFP)^57^ (**Fig. 7a, d**). Because the fluorescent channel previously used for unbiased cell membrane segmentation was now occupied by the ErkKTR-iRFP, we used the mEGFP channel to segment the cell outlines. To validate this approach, we used it to re-quantify our PMA-stimulation experiments and found that segmentation on the mEGFP channel leads to comparable results as when the myr-iRFP channel is used (**Fig. S10b**).

We simultaneously measured ADAM17 and ERK biosensor responses using the FR N/M ratio for eNRGies quantification and the ErkKTR-iRFP cytoplasm-to-nucleus (C/N) fluorescence ratio for ERK activity quantification. EGF treatment induced potent and rapid ERK activation, followed by an increase in ADAM17 activity that peaked at ∼100 min after EGF stimulation. As in the case of PMA stimulation, the EGF-induced ADAM17 response was blocked by inhibitors of ADAM17 (TMI), γ-secretase (DAPT), or the EGFR-ERK pathway (gefitinib and the MEK inhibitor PD0325901) (**Fig. 7e, f**, **Fig****. S10c, d, Movie S15**). ERK activity was abolished by EGFR/ERK pathway inhibitors, but not by protease inhibitors (**Fig. 7e, f**). Our biosensor thus confirms that ADAM17 is potently activated in MCF10A cells upon EGF-induced ERK stimulation, in agreement with studies in other epithelial contexts^20, 21, 23, 54^.

However, unlike PMA, EGF stimulation produced ADAM17 responses that were dynamic and variable between individual MCF10A cells, with some cells undergoing multiple clear pulses of nuclear accumulation (**Fig. 7f, Movie S16**). To quantitatively capture these single-cell dynamics, we segmented and tracked nuclei and cell bodies using the NLS-BFP and eNRGies mEGFP channels and quantified activity from each biosensor in single cells (see Methods). We also quantified NLS-BFP nuclear fluorescence in single cells as a control signal that should not be affected by stimuli. We processed trajectories by interpolating traces over small gaps and smoothing and subtracting slow drift over time before performing peak detection of activity pulses (**Fig. 7g, Fig. S11a, b**). Overall, our analysis produced biosensor responses in ∼750 cells that could be tracked throughout our ∼10 h experiments (**Fig. 7h**). Like ERK, ADAM17 activity was highly dynamic in single cells after EGF stimulation: most cells exhibited a high-amplitude pulse of ADAM17 activity at ∼100 min that fell back to baseline by ∼200 min, followed by asynchronous pulses with variable timing (**Fig. 7h**). These data demonstrate that eNRGies biosensors enable multiplexed detection of protease and signaling biosensor dynamics in individual living cells.

Finally, we set out to ask whether ADAM17 biosensor activity might be associated with prior or subsequent changes in ERK activity, or *vice versa*. We computationally aligned all cells’ ADAM17 pulses and measured the activity in the ErkKTR-iRFP and NLS-BFP channels from 150 min before and after the ADAM17 peak (**Fig. 7i, Fig. S11d, e**). We excluded the first peak identified in each trace to avoid capturing the acute biosensor responses driven directly by exogenous EGF stimulation (**Fig. 7h**). We also performed identical analyses for the ErkKTR-iRFP biosensor, where we aligned ErkKTR-iRFP activity pulses and measured ADAM17 and NLS-BFP responses before and after the ErkKTR-iRFP pulse (**Fig. 7i**, **Fig. S11d, e**). This analysis revealed that ADAM17 activity pulses are typically preceded by an ERK pulse by approximately 40 min; conversely ERK pulses are followed by an ADAM17 pulse after a ∼40 min delay. In contrast, the NLS-BFP nuclear fluorescence exhibited only minor fluctuations coinciding with the activity peaks, possibly due to a change in cell shape or volume or due to competition for shared nuclear transport machinery between all fluorescent components. We obtained similar results when only using the nuclear fluorescence intensity signals for our analysis (**Fig. S11a, e**).

In sum, our data reveals that ADAM17 activity is both ERK-dependent and dynamic after EGF stimulation in MCF10A cells. Multiplexed biosensor imaging further enables us to conclude that ERK and ADAM17 activity are dynamically interdependent. Computationally aligning pulses of either ERK or ADAM17 activity revealed corresponding pulses in the complementary biosensor separated by a time delay. Our data suggests that the relationship is asymmetric, with a pulse of ADAM17 activity following a pulse of ERK activity by approximately 40 min. Because ADAM17 is known to cleave EGF family members from the cell surface, potentially influencing ERK activity in neighboring cells, our data suggests that cycles of matrix metalloprotease cleavage may indeed play a role in the tissue-scale signaling dynamics observed for the EGF/ERK pathway.

## Discussion

Here we report the development of eNRGies: genetically encoded, modular biosensors for extracellular protease activity. Our biosensors are based on the principle of translating an extracellular peptide cleavage event into the membrane-to-nucleus translocation of an intracellular fluorescent protein. This process, termed regulated intramembrane proteolysis, is fundamental to how cells naturally relay extracellular cleavage events into the cell. eNRGies are also closely related to engineered γ-secretase-regulated proteins that have found widespread utility for mammalian synthetic biology, including the SynNotch and SNIPR systems^58, 59^. A key distinction is that our biosensor produces a rapid change in subcellular fluorescence, reporting on protease activity with dynamics on a timescale of tens of minutes, rather than a slower transcriptional response.

We show that the design is highly modular, working robustly for multiple cleavage sequences that can respond to a variety of extracellular and cell-surface proteases. It is also compatible with the release of multiple extracellular domains or intracellular reporters. Applying this biosensor to the ADAM17 cell-surface protease biosensor, we discover that ADAM17 is transiently activated during cell division in HEK293T cells and that ADAM17 is dynamically regulated in EGF-stimulated MCF10A cells in a manner that correlates with pulsatile ERK activity in individual cells. Overall, our data shows that the eNRGies scaffold enables the study of extracellular protease activity with high temporal resolution in individual living cells.

We systematically varied and/or optimized all components of the eNRGies scaffold: the bulky extracellular domain, extracellular linker, transmembrane domain, and intracellular output domain. Our experiments revealed a key design criterion of such a biosensor: the extracellular cleavage site must be separated by a long enough linker sequence from the transmembrane domain for good protease accessibility, but a short enough linker to promote secondary cleavage by the γ-secretase intramembrane protease. Our data suggests that these factors can be ideally balanced by linkers of ∼10-25 AAs, because we observed reduction in γ-secretase-dependent nuclear FR accumulation for longer of more structured linkers^29–31^. Moreover, the optimal linker sequence likely depends on the size of the extracellular protease domain and its proximity to the membrane. For example, we found that a linker of just 10 AAs performed well in the ADAM17 eNRGies system, matching the cleavage site architecture of endogenous substrates. These data suggest that the distance of the cleavage site from the membrane of natural substrates of cell-surface proteases should be considered in the biosensor design. It is notable that some natural and synthetic γ-secretase substrates have larger ECDs and yet are reported to be efficiently cleaved^60, 61^, a phenomenon that warrants further mechanistic study.

The eNRGies scaffold relies on the activity of γ-secretase for nuclear translocation of its fluorescent reporter. This requirement could in principle pose a challenge in cells where γ-secretase is not present. Fortunately, γ-secretase activity is thought to be ubiquitous: it is conserved in metazoans^29^ and is expressed across mice and human tissues, and cell lines^62, 63^. The level of γ-secretase expression in HEK293T cells is among the lowest reported in the Human Protein Atlas^63^, yet even in HEK293Ts, extracellular domain cleavage is followed within minutes by γ-secretase-dependent nuclear translocation of the intracellular fluorescent protein (**Fig. 2g**). These data suggest that even low levels of γ-secretase are sufficient for robust biosensor function. Finally, we note that the biosensor’s dependence on γ-secretase activity also enables blockade of biosensor responses using pharmacological inhibitors such as DAPT as a negative control or potentially to synchronize biosensor responses by inhibitor washout.

We define two strategies for biosensor quantification based on analysis of the translocation from membrane to nucleus of a fluorescent intracellular domain. First, we show that the nuclear-to-membrane ratio serves as an excellent metric for protease activity, responding robustly and sensitively across cellular contexts and proteases. Because a single fluorescent protein is quantified ratiometrically, our measurement is independent of overall fluorescent protein expression level, permitting reproducible quantification across experiments and target proteases. The nuclear-to-membrane ratio is dominated by the nuclear signal, so that it can yield reliable signal even with imperfect membrane segmentation (e.g. using the biosensor fluorophores) (**Fig. 7**). We find that nuclear fluorescence alone, apart from delivering a qualitative, visual readout, can also provide an excellent readout of protease dynamics albeit in a manner that still depends on each cell’s biosensor expression level (**Fig. 2, Fig. S11**). This suggests that eNRGies can provide an excellent readout of protease dynamics even where membrane fluorescence is difficult to measure in individual cells, for example when cells are in close contact in tissues (**Fig. 7**).

As a key application of the eNRGies platform, we developed a novel biosensor with high specificity for the ADAM17 cell-surface protease to enable high-resolution studies of its roles in cell signaling. We show that biosensor activity is abolished in cells knocked out for ADAM17 or treated with TMI-1, a small molecule ADAM17 inhibitor (**Fig. 5**).

Our biosensor enabled the immediate discovery of dynamic ADAM17 activity in live cells. We found that HEK293T cells transiently activated ADAM17 upon cell division, exhibiting an increase in nuclear biosensor accumulation after completion of mitosis that was not observed in ADAM17 knockout cells (**Fig. 6**). Our observation is consistent with two prior studies suggesting association between ADAM17 regulation and mitosis^64, 65^. While the mechanistic basis for mitosis-coupled ADAM17 activation is unclear, ADAM17 activity has recently been shown to be regulated by phosphorylation of a regulator, iRhom2, by the MAPKs ERK, JNK and p38 on its intracellular domain^66, 67^. It may also be the case that iRhom2 is further regulated by G2 cyclin/CDK activity (which, like the MAPK protein kinase family, phosphorylates Ser/Thr-Pro peptide sequences).

Applying the biosensor in MCF10A cells further revealed that ADAM17 activity is dynamically regulated following epidermal growth factor (EGF) stimulation. EGF stimulation produces a high-amplitude transient pulse of ADAM17 activity. Following this initial activation event, we observe spontaneous pulses in both ADAM17 and ERK activity, with ADAM17 pulses correlated with a preceding pulse of ERK activity in the same cell ∼40 min earlier (**Fig. 7**). Our biosensor thus reveals that ADAM17 activity is variable on a timescale of tens of minutes, a timescale that is relevant to the complex dynamics of EGF/ERK activity that is often observed in epithelial monolayers (typically 0.5-2 pulses per hour)^68, 69^. It will be important in future studies to perform single-cell analysis in spatial neighborhoods to test whether stochastic pulses of protease activity produce paracrine responses. Such an analysis could potentially link spatiotemporal dynamics of ADAM17 to the propagation of ERK signaling waves and pulses observed during wound healing, local oncogene activation, apoptosis or infection^52, 53, 70–74^. Analyzing ERK and ADAM17 activity as done here under mechanical perturbations may also help to further explain the interplay between the EGFR-ERK pathway and ADAM17 activity^22, 45, 72^.

Beyond the applications demonstrated here, we envision that the eNRGies biosensor platform could find diverse applications for drug discovery and synthetic biology. eNRGies could be combined with pharmacological inhibitors or genetic perturbations to drive new insights into substrate repertoire, protease redundancy, or to identify new small-molecular protease inhibitors. For example, cells expressing eNRGies with libraries of cleavage peptide sequences could be analyzed in imaging-based screens to map the specificity of an extracellular protease of interest. eNRGies-expressing cells could be treated with candidate small molecules, fixed, and assessed for nuclear fluorescence to identify candidate protease inhibitors. Our biosensor platform could also be further functionalized by replacing the intracellular fluorescent protein with an effector domain (e.g., a transcription factor) to “wire in” extracellular protease activity to novel cellular responses for immunotherapy or other applications.^75^

## Materials and Methods

### Plasmids

For cloning all plasmids were propagated in *E. coli* Stellar Competent Cells (Takara, #636763). We used In-Fusion seamless assembly for the cloning of larger DNA fragments into vectors. DNA fragments and vector backbones were amplified by PCR with primers that contained 15-20 bp overlaps between adjacent fragments. The PCR products were then combined using In-Fusion Snap Assembly Master Mix (Takara, #638947) according to manufacturer’s instruction. We used blunt end cloning for the introduction of shorter peptide sequences. The entire vector was amplified via PCR with primers containing the additional sequences and subsequently ligated using the KLD Enzyme mix according to the manufacturer’s instruction (NEB, #M0554S). All constructs were cloned into a piggyBac vector (System Biosciences) with a SFFV promoter. pQC110 was made by Qinhao Cao^76^. The CD28 sequence for the plasmid pBR20 was amplified from pEF6a-FKBP-CD28 which was a gift from Min Zhuang (Addgene plasmid # 113401)^77^. The Halotag sequence for the plasmid pBR252 was amplified from pJL3 which was a gift from Josh Levitz. A list of all plasmids and protein sequences used in this study can be found in Table S1 and Table S2, respectively.

### Cell line generation

Clonal transgenic cell lines were generated via transfection and PiggyBac genomic integration. Cells were plated 24 h before transfection in a 12 well plate. 830 ng of the target PiggyBac vector plasmid and 185 ng of the PiggyBac transposase plasmid were transfected using Lipofectamine 3000 Transfection Reagent (Invitrogen, #L3000015) according to manufacturer’s instruction. HEK293T cell lines were passaged the next day. For MCF10A-5E cell lines^78^, the medium was replaced with fresh medium four hours after transfection and cells were passaged two days later. Four to six days after transfection, cells positive for the respective fluorescent protein marker were sorted by fluorescence-activated cell sorting (Sony, SH800S). For HEK293T cell lines, monoclonal lines for the membrane and nuclear marker were generated. All subsequent biosensor HEK293T cell lines were polyclonal unless otherwise noted. For MCF10A-5E, monoclonal cell lines were generated. A list of all cell lines used in this study can be found in Table S3.

### Cell culture

HEK293T cell lines were cultured in high-glucose DMEM (Gibco, #11995073) supplemented with 10% fetal bovine serum (R&D Systems) and 100 units/ml penicillin/streptomycin (Gibco, #15140122). For imaging HEK293T cells were maintained in OptiMEM (Gibco, #31985070). MCF10A-5E cell lines^78^ were cultured in DMEM/F12 with HEPES (Gibco, #11320032) supplemented with 5% horse serum (Millipore Sigma, #H1138), 20 ng/mL EGF (R&D Systems, # 236-EG), 0.5 μg/mL hydrocortisone (Millipore Sigma , #H-0888), 100 ng/mL cholera toxin (Millipore Sigma, #C8052), 10 μg/mL insulin (Millipore Sigma, #I1882), and 100 units/ml penicillin/streptomycin (Gibco, #15140122). MCF10A starvation media consisted of phenol-red free DMEM/F-12 with HEPES (Gibco, #11039021) supplemented with 0.5% horse serum and 1 mM Na pyruvate (Gibco, # 11360070). All cells were maintained at 37°C and 5% CO2. Cells were tested to confirm the absence of mycoplasma contamination using a PCR-based universal Mycoplasma detection kit (ATCC, #30-1012K).

### Preparation of cells for live-cell imaging

Cells were plated on fibronectin-coated, 96-well plates with a #1.5 cover glass bottom (Cellvis, #P96-1.5H-N). Plates were coated with 10 μg/ml fibronectin diluted in PBS for 30 min. HEK293T cells were plated in 200 μl full media at a density of 60000-70000 cells/well and MCF10A cells were plated in 200 μl full media at a density of 5000-7000 cells/well 18 to 24 h before imaging. Sixty minutes before image acquisition, the medium was exchanged to 200 µl OptiMEM for HEK293T cells or to starvation medium for MCF10A cells. Inhibitors were added 60 min before image acquisition.

### HaloTag labeling

Cells expressing a HaloTag labeled biosensor were incubated with 100 nM Janelia Fluor JFX554 HaloTag ligand (Promega, # HT1030) for 1 h in 200 µl full media before the medium was exchanged to 200 µl OptiMEM without additional washes for image acquisition.

### Live-cell imaging

Timelapse imaging of cells was performed on a Nikon Eclipse Ti microscope equipped with a Yokogawa CSU-X1 spinning disk, an Agilent laser module containing 405, 488, 561, and 650 nm lasers, and an iXon DU897 EMCCD camera. All images were acquired using 40× or 60× oil objectives. Cells were maintained at 37°C with 5% CO2 during image acquisition by an environmental control unit (Okolab). Additionally for longer time-series (> 3 h) cells were covered with mineral oil (Millipore Sigma, # 330779) to prevent evaporation of media.

### Recombinant proteases

TEV protease (NEB, #P8112S) was used at 5/10/20 Units, Enterokinase light chain (NEB, #P8070S) at 32 Units, Factor Xa Protease (NEB, #P8010S) was used at 1 µg and active, human, recombinant MMP-9 (Millipore Sigma, #PF140) used at 0.2 µg corresponding 27.6 Units per well of a 96-well plate in 200 µl media.

### Inhibitory and stimulating agents

Cells were preincubated with inhibitory agents 1 h prior to image acquisition. Inhibitory agents were used at the following concentrations per well of a 96-well plate in 200 µl OptiMEM or starvation media for HEK293T cells and MCF10A cells, respectively: MEK inhibitor MEK1/2 inhibitor PD 0325901 (Tocris, 4192), #SML0789), EGFR inhibitor Gefitinib (MedChemExpress, #HY-50895) at 5 µM, p38 MAPK inhibitor SB202190 (Selleckchem, # S1076) at 1 µM, γ-secretase inhibitor DAPT (Millipore Sigma, #D5942) at 10 μM, and γ-Secretase Inhibitor XXI, Compound E (Millipore Sigma, #565790) at 100 nM, ADADM17 inhibitor TMI-1 (MedChemExpress, #HY101448) at 10 µM.

Addition of stimulating agents was performed during image acquisition unless otherwise noted and the first frame after addition was defined as t = 0. Stimulating agents were used at the following concentrations per well of a 96-well plate in 200 µl OptiMEM or starvation media for HEK293T cells and MCF10A cells, respectively: 100 nM PMA (Millipore Sigma, #524400), 10 µg/ml anisomycin (Sigma-Aldrich, #A9789) and 100 ng/ml EGF (R&D Systems, #236-EG).

### Image analysis

The processing, analysis and visualization of all data was performed using custom-written scripts in Jupyter Notebooks with Python 3.11.15 and the associated packages numpy (2.0.2), pandas (3.0.1), scipy (1.17.1), matplotlib (3.10.8), seaborn (0.13.2), tifffile (2026.3.3), IPython (9.10.1) and jupyter_core (5.9.1).

Cell bodies and nuclei were segmented using Cellpose^38^. All HEK293T data was segmented using Cellpose2^79^: the nuclear, NLS-BFP channel was segmented using the model “nuclei” and a radius of 48; the cell body, myr-iRFP channel was segmented using the model “cyto2” with a radius of 65 and both the myr-iRFP and nuclei channel as input. All MCF10A image data was segmented using cellpose 3^80^: the nuclear, NLS-BFP channel was segmented using the model “nuclei”, the restore type onclick_nuclei and a radius of 40; the cell body, myr-iRFP channel was segmented using the model “cyto3”, the restore type oneclick_cyto3 and a radius of 60 and both the myr-iRFP and nuclei channel as input. For MCF10A that harbored an ErkKTR-iRFP instead of the myr-iRFP the mEGFP channel was used for cell body segmentation with the same parameters as described.

The cellpose segmentation outputs cell body and nuclear masks in which each cell or nucleus is associated with a labeled ROI with increasing gray value, and a binary cell outline mask. We modified the cellpose python scripts to generate binary masks from the cell body and nuclear masks for field-of-view level analysis. For analysis of the ErkKTR-iRFP data we further used the nuclear mask to generate an 8-pixel wide cytoplasm ring mask around the nucleus.

Using custom-written python scripts, we then extracted fluorescence intensities and fluorescence intensity concentrations. All reported intensity and intensity ratio values are fluorescence intensity concentrations or mean fluorescence intensity per pixel. The following masks were used to extract the respective intensities: cellpose nuclear masks were used for nuclear fluorescence intensity, cellpose cell outline masks were used for membrane fluorescence intensity, and both were used to calculate the nuclear-to-membrane (N/M) fluorescence ratio. For ErkKTR-iRFP activity we used the cytoplasm ring mask for cytoplasm intensity and the nuclear mask for nuclear intensity, and both were used to calculate the cytoplasm-to-nucleus (C/N) fluorescence ratio.

Three distinct analysis modes were used: In the field of view level analysis, fluorescence intensities and ratios were measured for the whole field of view using the binary nuclear masks and cell outlines. The size of the field of view analyzed for HEK293T cells was 230.45 µm x 230.45 µm and for MCF10A 346.38 µm x 346.38 µm. In the single-cell mode in which cells are not tracked, fluorescence intensities and ratios were measured for individual cells by using the cellpose cell body masks. Note that the gray value associated with each ROI in the cellpose cell body and nuclei masks are not consistent across frames, nor are they associated with each other. To link membrane and nuclear fluorescence intensity we used the cellpose cell body mask and the binary nuclear and cell outline masks to generate nuclear and membrane masks with a unique ID per field of view linked to the gray value of the ROI of the cell body mask. In a third scenario in which single cells are tracked across frames we used cell body masks where the ID (gray value associated with the ROI) for a tracked cell persists across frames, which were then combined with the binary nuclear, cell outline and cytoplasm ring masks to extract intensities for a given cell across frames.

### Single-cell tracking

For the tracking of HEK293T cells through the division, we used the fiji (version 1.54p) plugin trackmate (version 7.14.0)^81^. For tracking we used the cellpose cell body mask with the label image detector and the LAP tracker (linking max distance 20 µm; track splitting; all with a penalty of 1-3 on the median intensity of the NLS-BFP fluorescence channel). Tracks were filtered to allow for less than 1.7 split events and minimum duration of 22 frames. The resulting label image with cell body masks with persistent ID and the spots.csv containing the spot (object/cell) ID and track IDs were saved for downstream analysis.

For the tracking of MCF10A, cells exposed to EGF cells were tracked using a custom-written python script with a nearest-neighbor linking algorithm. Nuclear centroids were extracted from the cellpose nuclear masks for each frame and linked across frames by solving a linear assignment problem (Hungarian algorithm, scipy.optimize.linear_sum_assignment) on the matrix of Euclidean centroid distances, following the linear assignment problem (LAP) framework^82^ with the following parameters: linking max distance of 20 µm, track segment gap closing of 15 µm and 2 frames; minimum track length of 40 frames. For each tracked nucleus, the corresponding cell body was assigned per frame as the cell-body label from the cellpose cell body masks with the greatest pixel overlap with the nucleus. The resulting cell tracks were written as a label image with cell body masks with persistent ID, together with a spots.csv file matching the TrackMate output format for downstream analysis.

### Alignment of cell division events in HEK293T single cell tracks

Single-cell fluorescence intensity ratio tracks were aligned to mitosis time = t0 defined as the first occurrence of two cells (spots) with the same track ID in the trackmate file. Only those cells whose tracks fully spanned the analysis window were retained.

### Peak detection and peak-triggered averaging of eNRGies and ErkKTR-iRFP pulses in EGF-stimulated MCF10A

To understand whether eNRGies and ErkKTR-iRFP biosensor are temporally correlated, we calculated peak-triggered averages. Since the nuclear fluorescence intensity of the ErkKTR-iRFP decreases upon ERK activation as the reporter translocates from the nucleus into the cytoplasm, the iRFP nuclear fluorescence intensity was sign-inverted prior to peak detection and averaging so that increases reflect reporter activity. The nuclear fluorescence intensity of the nuclear marker NLS-BFP was used as a control that should not respond to EGF. For each tracked cell, single-cell fluorescence intensity and ratio traces were then baseline-corrected and smoothed: a rolling baseline was estimated as the 50th percentile over a 41-frame window and subtracted, followed by Gaussian smoothing (*σ* = 2 frames). Tracks were only retained if they started within the first 5 frames, i.e. before EGF addition, and covered more than 90% of the analysis window. Gaps from missed detections/segmentation of the nucleus or cell body, outlier removal for the fluorescence intensity ratios (absolute ratio > 5), and zero values were filled by piecewise cubic Hermite interpolation. Pulses were detected on the processed traces using `scipy.signal.find_peaks` with a prominence threshold scaled to each cell’s signal range (0.3 for the FR and NLS-BFP N/M ratios, 0.2 for the iRFP and NLS-BFP C/N ratios and iRFP, FR and NLS-BFP nuclear intensities), a minimum peak separation of 5 frames, and a minimum width of 3 frames.

To examine the temporal relationship between biosensor signals, peak-triggered averages were computed. For each detected peak (excluding each cell’s first peak), a window spanning 30 frames before and 30 frames after the peak was extracted from all co-recorded signals and aligned to the peak time (t = 0). For the fluorescence intensity ratio readouts, the extracted windows were averaged directly. For the nuclear intensity readouts, each window was first baseline-subtracted using the mean of its pre-peak interval before averaging, to account for the differing absolute scales of the raw intensities.

### Fitting of fluorescence intensity time courses

Decay of mEGFP and FR fluorescence on the membrane after TEVp exposure, and decay of the FR N/M ratio and FR fluorescence in the nucleus after TEVp washout were fitted to a delayed exponential decay model, defined as *y*(*t*) = *C* for *t* < *t*_0_ and 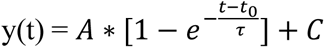 for *t* ≥ *t*_0_. Increase of FR fluorescence in the nucleus and the rise of the FR N/M ratio after TEVp exposure after TEVp washout were fitted to a delayed exponential decay model, defined as *y*(*t*) = *A* + *C* for *t* < *t*_0_ and 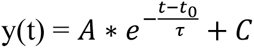 for *t* ≥ *t*_0._ The increase of mEGFP fluorescence on the membrane after TEVp washout was fitted to an exponential rise model, defined as 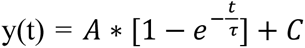. The free parameters *A*, *τ*, *t*_0_ and C, where *A* is the amplitude, *τ* is the time constant, *t*_0_ the onset delay and *C* is the offset. The initial fit parameters were estimated in the following way: *A*_0_= max(*y*) – min(*y*); *τ*_0_= 0.5 ∗ (max(*t*) – min(*t*)), *C*_0_= min(*y*) and *t*_0_ = min(*t*). For fits after TEVp addition, the fit was performed between 0 and 55 min, except for the FR fluorescence in the nucleus after exposure with 20 U TEVp, where the fit was performed between 0 and 25 min. For fits after TEVp washout, the fit was performed between 60 min and 760 min after TEVp addition.

The decay of the FR N/M ratio and FR fluorescence in the nucleus after TEVp washout was fitted with the same function. The initial fit parameters were m0 = 0.7; *τ*0 = 120; b0 = 0.1 and the fit was performed between 80 min and 760 min post TEVp addition. *τ* was subsequently used to calculate the half-life time 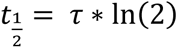. The increase in mEGFP fluorescence on the membrane was fitted to the function 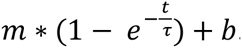. The fit is performed with the same initial parameters between 80 min and 465 min post TEVp addition. *τ* was subsequently used to calculate the half-rise time 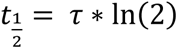.

## Data availability

The raw microscopy images that support the findings of this study are available from the corresponding authors upon request. Major plasmids associated with this study are being deposited on Addgene (plasmids #259190-259203).

## Code availability

Example code for the analysis will be made available on GitHub, all other code is available from the corresponding authors upon request.

## Acknowledgements

We thank Andre Frankenthal, John Ngo, and members of the Toettcher lab, particularly Qinhao Cao and Harrison Oatman, for helpful discussions. We thank Veit Hornung (LMU) for providing the HEK293T ADAM10/ADAM17 KO cell lines. This work was supported by the NSF Center for the Physics of Biological Function PHY-1734030 (to B.R.), the Max Planck Society (to B.R.) and by the National Institutes of Health grant R01GM144362 and R35164185 (to J.E.T.).

## Author contributions

Conceptualization, B.R. and J.E.T.; Investigation, B.R. and M.W.; Writing – Original Draft, B.R.; Funding, B.R. and J.E.T.; Writing – Review & Editing, B.R. and J.E.T

