## Supplementary Information for "A modular live-cell biosensor for extracellular protease activity"

**Supplementary Figures S1-S11**

**Supplementary Movie Legends Movie S1-S16**

**Supplementary Tables S1-S3**

### Supplementary Figures

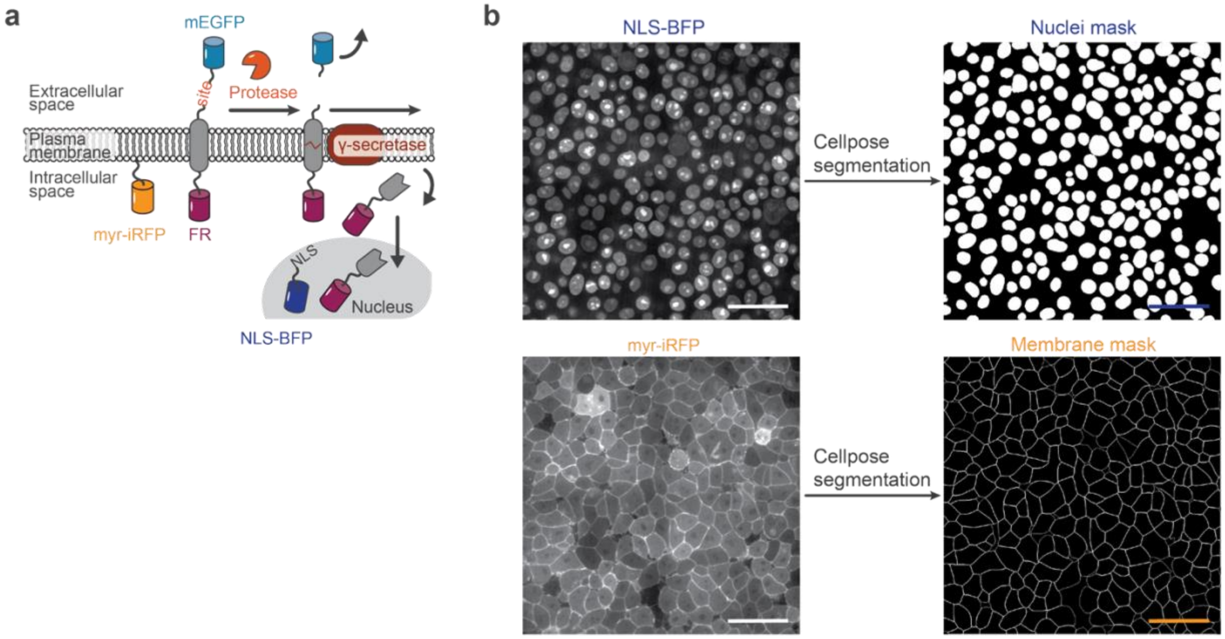


**Fig. S1: Segmentation of membrane and nuclei markers for unbiased quantification of membrane and nuclear fluorescence intensities of the protease biosensors. a,** Schematic of the biosensor mechanism and nuclear and membrane markers (NLS-BFP and myr-iRFP, respectively) used for segmentation. **b,** Monoclonal cell lines express NLS-BFP and myr-iRFP as nuclear and membrane markers, respectively, and the respective fluorescent channels are acquired for every timepoint and subsequently used to segment the nuclei and the membranes using Cellpose (see methods for details). Scale bars, 50 µm.

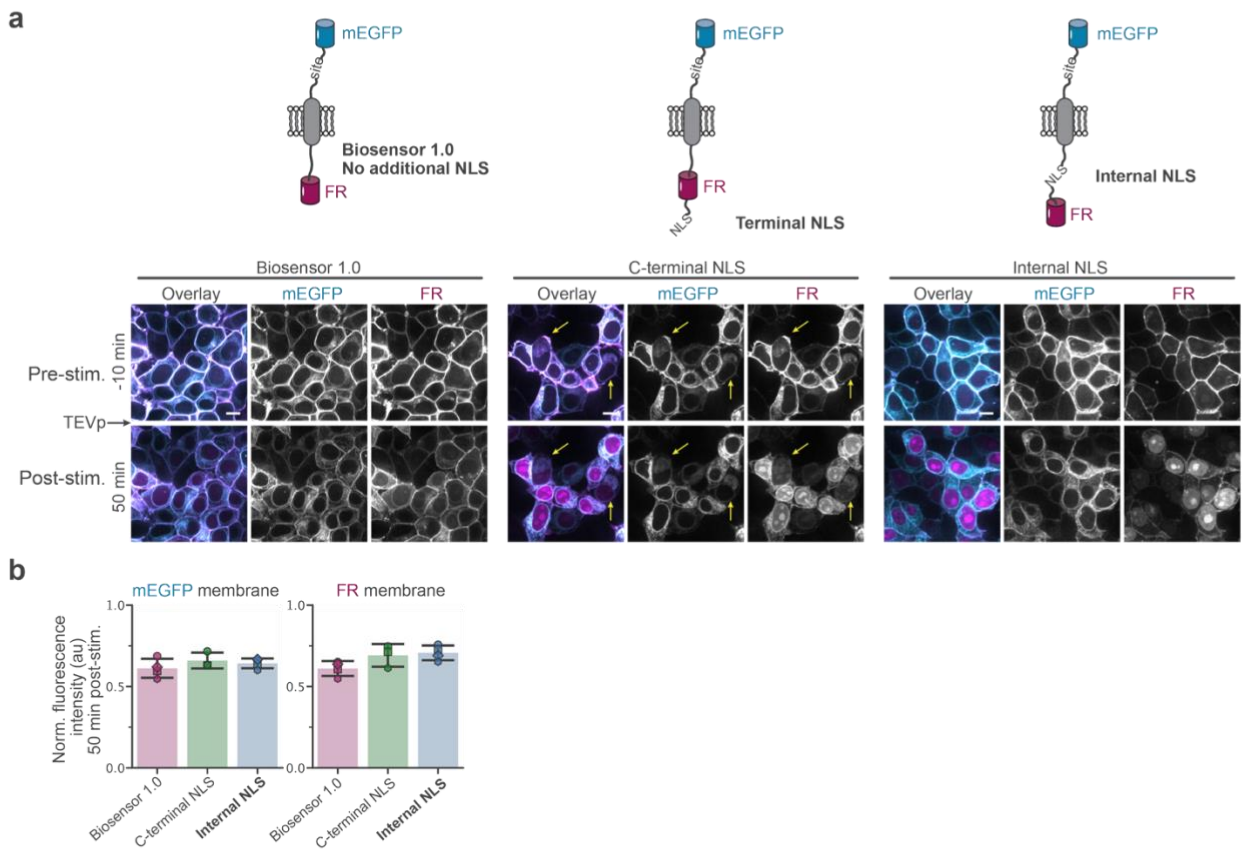


**Fig. S2: Biosensors with additional NLS show nuclear localization of the ICD upon TEVp exposure, but a C-terminally located NLS leads to occasional mis-trafficking. a**, Representative images of cells expressing the initial design (biosensor 1.0) that has no additional NLS, or designs with an additional NLS at the C-terminal end (C-terminal NLS) or after the juxtamembrane domain (internal NLS) before and 50 min after exposure to 20 U TEVp. Yellow arrows highlight cells in which the biosensor has not trafficked to the cell surface, but seems primarily localized to the ER/Golgi. **b**, Normalized fluorescence intensity of mEGFP and FR on the membrane 46-54 min after TEVp exposure for the original design and designs incorporating an additional NLS directly after the juxtamembrane domain or at the C-terminal end. Bar and error bars represent the mean ± SD from *N*(Biosensor 1.0) = 4, *N*(C-terminal NLS) = 3, *N*(Internal NLS) = 4 independent experiments identified by marker, respectively. Scale bars, 10 µm.

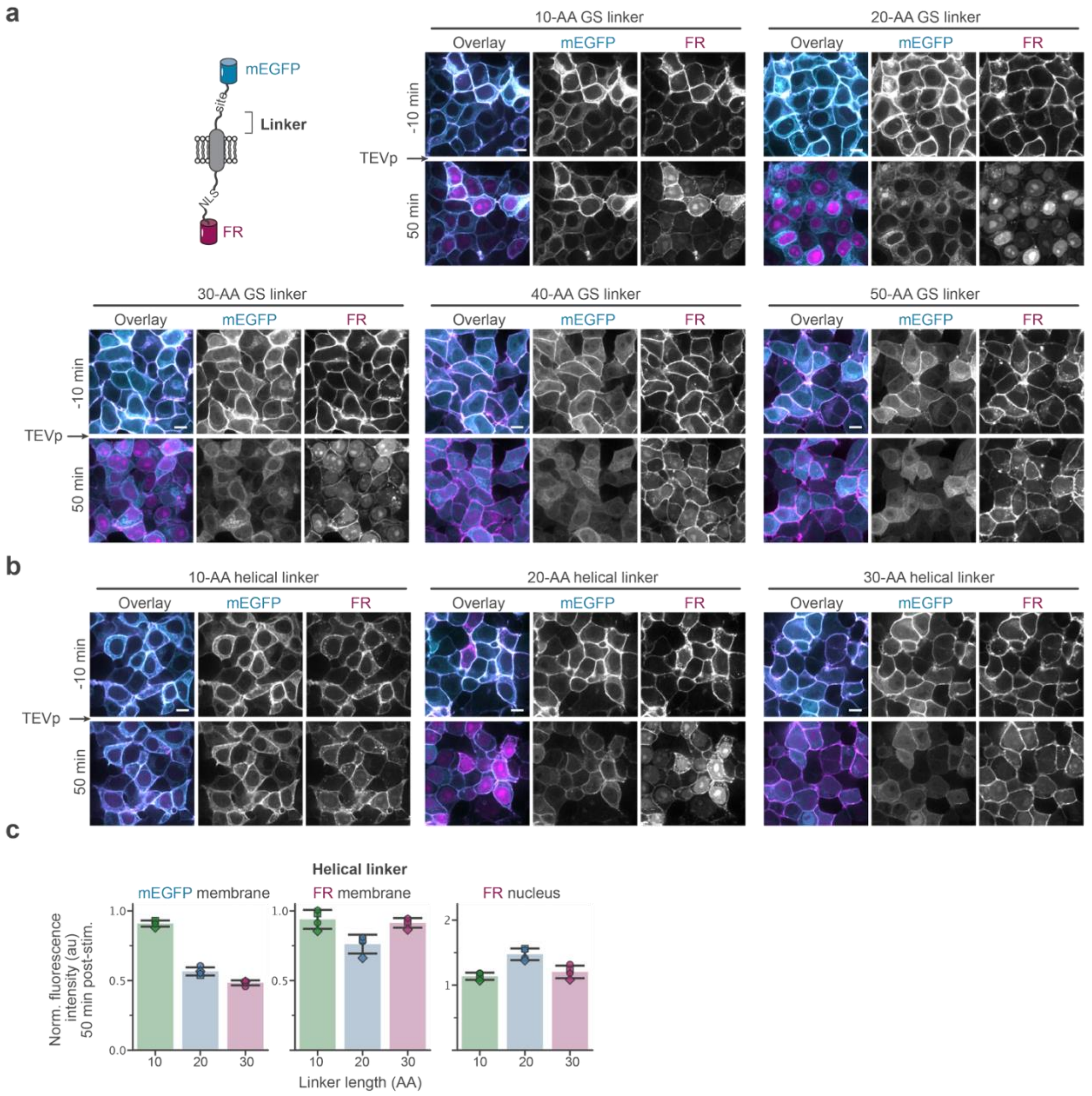


**Fig. S3: Biosensors with a helical linker are not efficiently cleaved by γ-secretase. a**, Representative images of cells expressing a biosensor with additional internal NLS in which the original NRG linker sequence has been replaced with a GS linker of varying length (10-50 AA) before and 50 min after exposure to 20 U TEVp. **b**, Representative images of cells expressing a biosensor with additional internal NLS in which the original NRG linker sequence has been replaced with a helical linker of varying length (10, 20 and 30 AA) before and 50 min after exposure to 20 U TEVp. **c**, Normalized fluorescence intensity of mEGFP and FR on the membrane and FR in the nucleus 46-54 min after TEVp exposure for biosensors with additional internal NLS and a helical linker of varying length (10, 20 and 30 AA). Bar and error bars represent the mean ± SD from 4 independent experiments identified by marker, respectively. Scale bars, 10 µm.

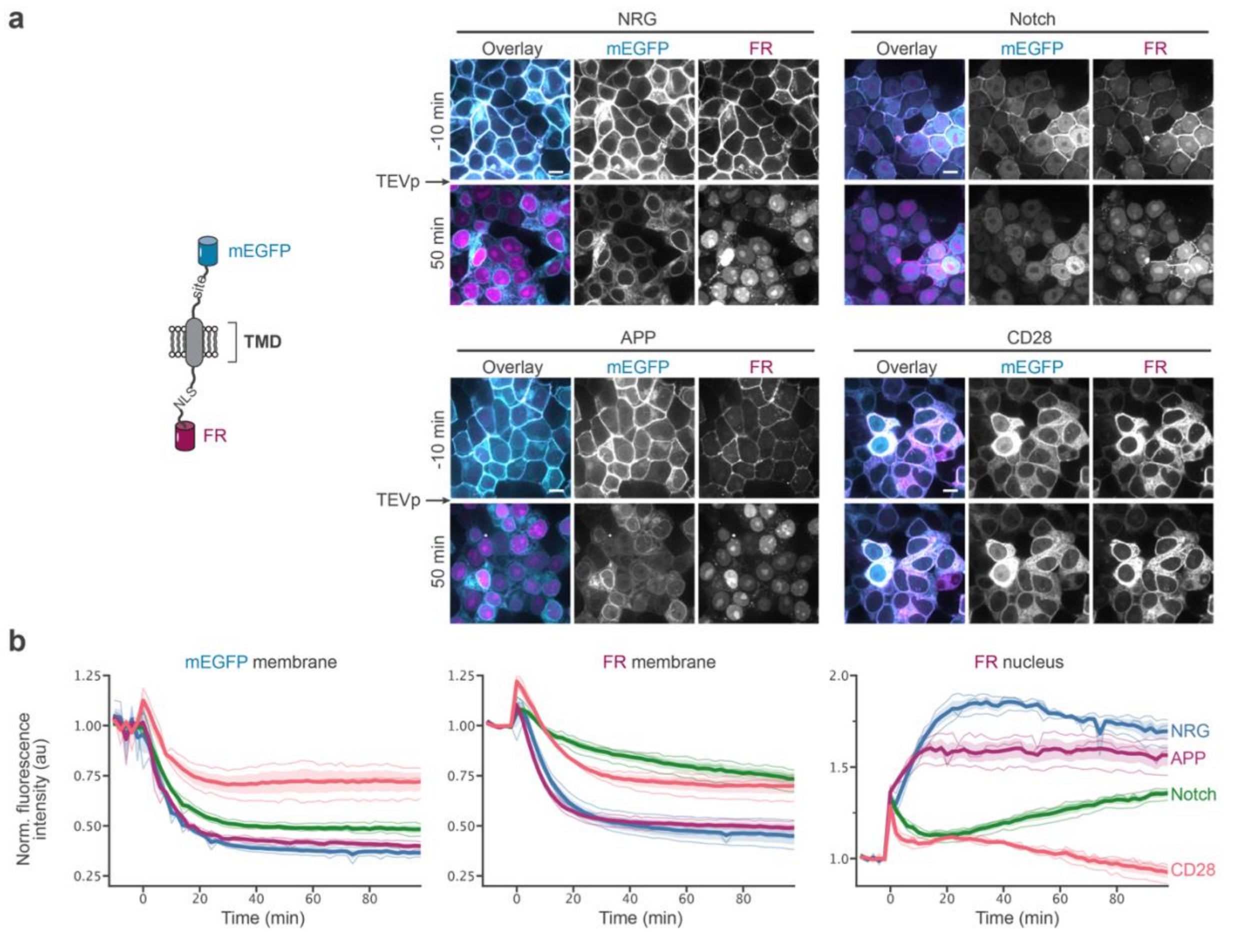


**Fig. S4: Biosensor with the NRG transmembrane domain outperforms those with an APP, Notch or CD28 transmembrane domain. a,** Biosensors incorporating APP, Notch or CD28 as transmembrane domain are partially mis-localized to the nucleus or ER. Representative images of cells expressing the optimized biosensor with different transmembrane domains that are known γ-secretase substrates, NRG, Notch, APP or as control CD28 that is not a γ-secretase substrate. **b,** Normalized fluorescence intensity time courses of mEGFP and FR on the membrane and FR in the nucleus of cells expressing the biosensors with different transmembrane domains. Cells are exposed to 20 U TEVp at time = 0. The bold lines and band represent the mean ± SEM of traces from three independent experiments. Scale bars, 10 μm.

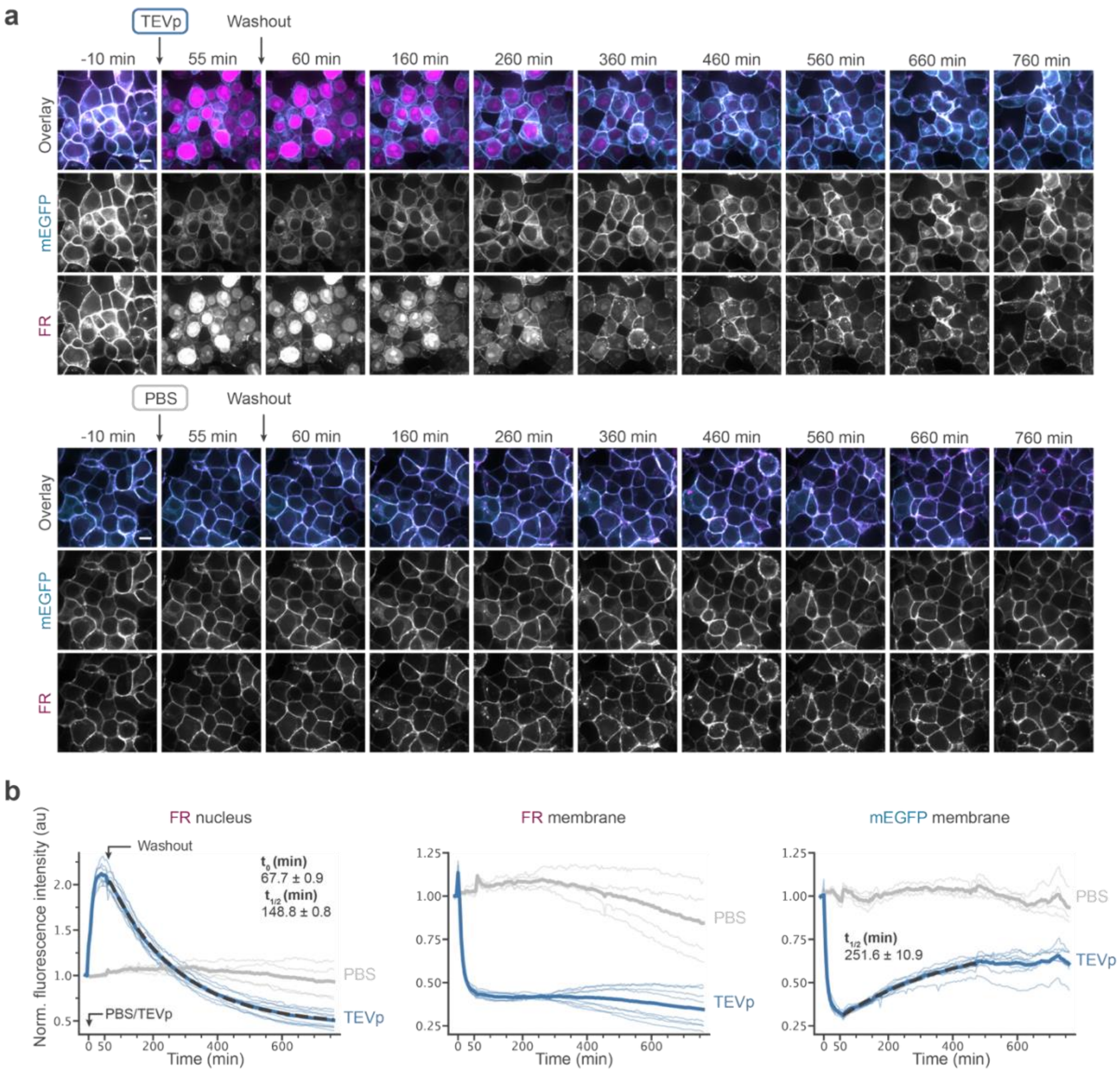


**Fig. S5: Recovery of the N/M ratio is dominated by recovery of the nuclear signal upon removal of the activating protease. a, b,** Representative images (**a**) and normalized fluorescence intensity time courses of mEGFP and FR on the membrane and FR in the nucleus (**b**) of cells expressing the TEVp biosensor that were exposed to PBS or 20 U TEVp at time = 0 and were washed after 55 min of incubation to remove PBS or TEVp. Bold lines represent the mean of traces from *N*(PBS) = 4, *N*(TEVp) = 8 independent experiments, respectively. (Delayed) exponential fits to the response are shown in dark dashed lines and the delay parameter and half-lives are indicated. Scale bars, 10 µm.

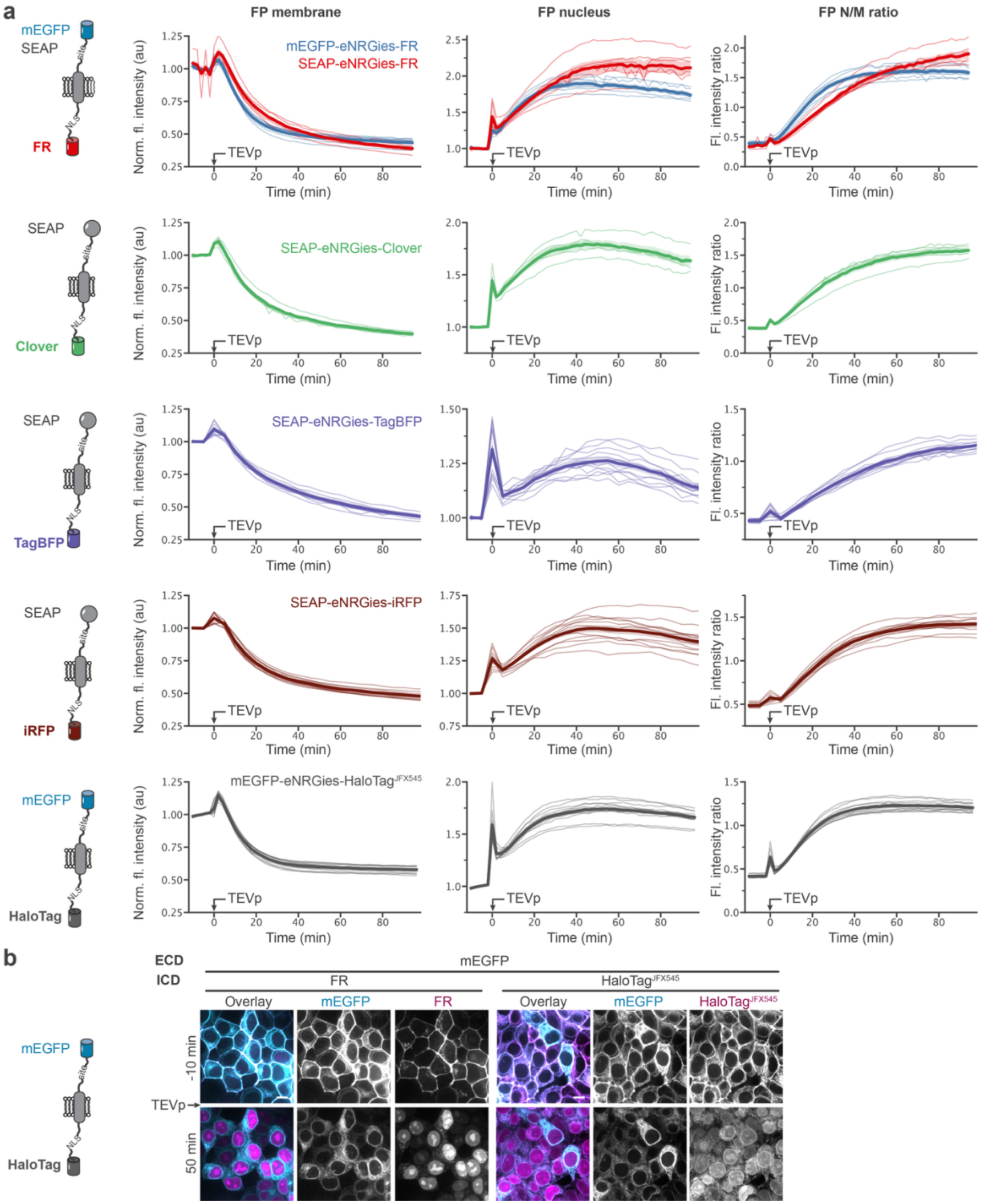


**Fig. S6: eNRGies are a modular biosensor platform across multiple ECDs and ICDs a,** Time course of the normalized fluorescence intensity of the ICD channel on the plasma membrane and the nucleus of cells expressing a TEVp eNRGies with varying ECD and ICDs. eNRGies with a HaloTag ICD were labeled with a JFX554 HaloTag ligand before image acquisition. Cells are exposed to 20 U TEVp at time = 0. The bold lines and band represent the mean ± SEM of traces from *N*(mEGFP-FR) = 6, *N*(SEAP-FR) = 6, *N*(SEAP-Clover) = 5, *N*(SEAP-BFP) = 12, *N*(SEAP-iRFP) = 12, *N*(mEGFP-HaloTag) = 12 samples and three independent experiments, respectively. **b**,

Representative images of cells expressing a TEVp eNRGies with mEGFP as ECD and FR or a HaloTag ICD labeled with the JFX545 HaloTag ligand before or 50 min after exposure to 20 U of TEVp at time = 0. Scale bars, 10 μm.

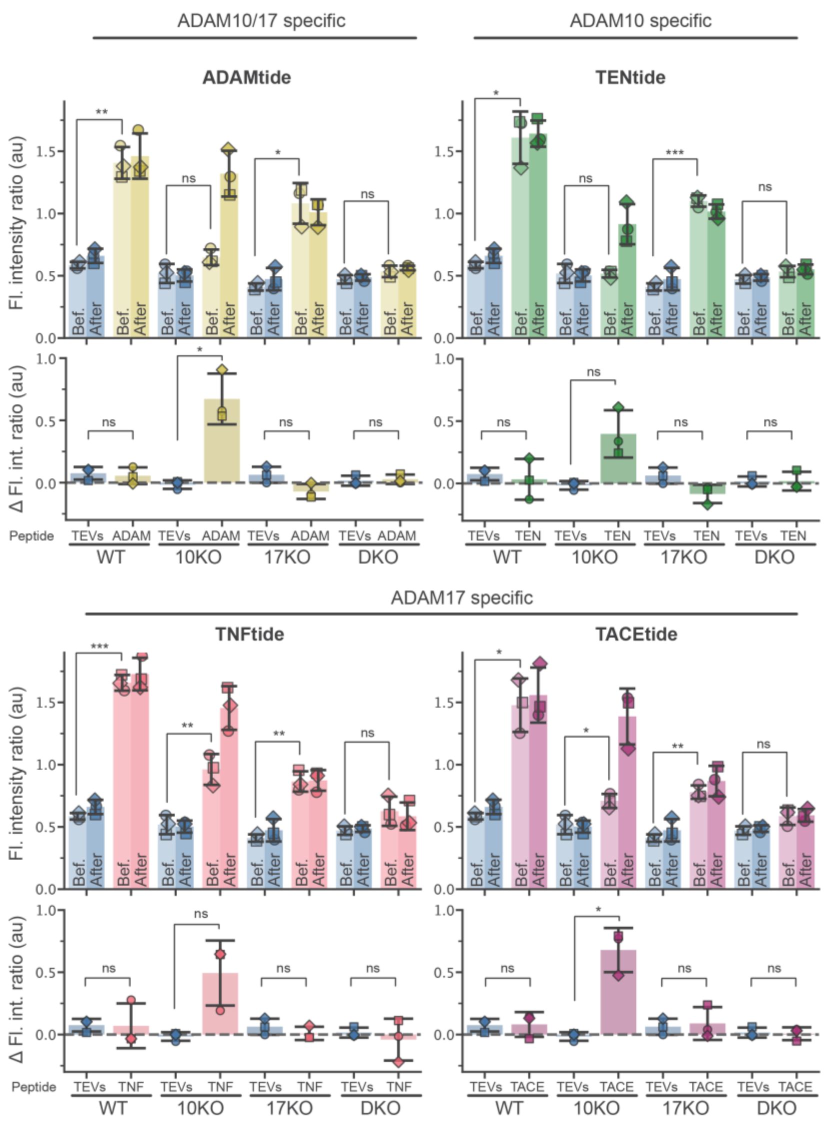


**Fig. S7: eNRGies with optimized peptide sequences are not suitable as ADAM10 or ADAM17 biosensors.** FR N/M ratio (top) and change in FR N/M ratio (bottom) of WT HEK293T (WT), ADAM10KO (10KO), ADAM17KO (17KO) and double ADAM10/ADAM17KO (DKO) cells expressing eNRGies with TEV-site (TEVs) as control, or eNRGies biosensors with the putative ADAM10 and ADAM17 substrates ADAMtide, TENtide, TNFtide and TACEtide before (-10 min) and after (70/80 min) stimulation with PMA. Bar and error bars represent the mean ± SD from three independent experiments identified by marker. Statistical analysis was conducted using a two-sided Welch's *t*-test. ns > 0.05, *$P \leq 0.05$, **$P \leq 0.01$ and ***$P \leq 0.001$.

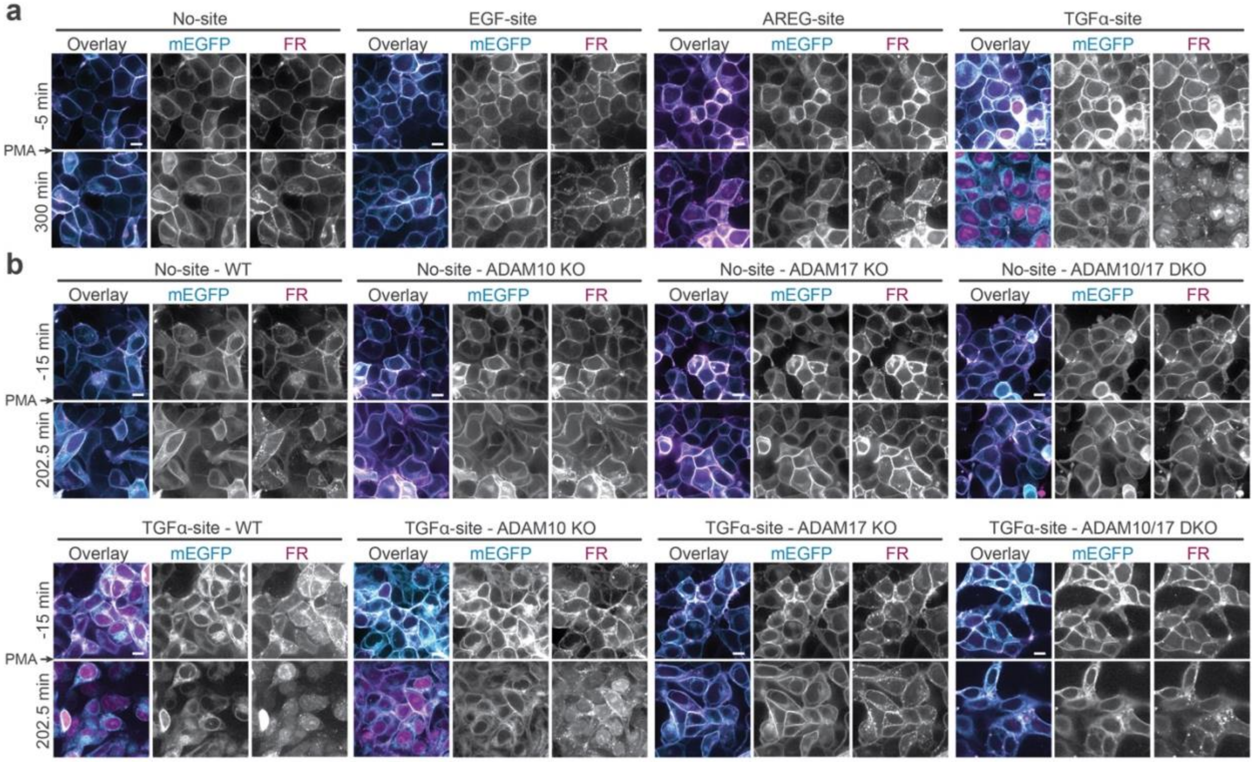


**Fig. S8: TGFα-site eNRGies reports on the activity of the sheddase ADAM17. a,** Representative fluorescence images of cells expressing a site-less eNRGies (No-site) or eNRGies with the proximal cleavage sites of EGF, AREG or TGFα before and after stimulation with PMA. **b**, Representative fluorescence images of WT HEK293T (WT), ADAM10KO (10KO), ADAM17KO (17KO) and double ADAM10/ADAM17KO (DKO) cells expressing eNRGies with the proximal cleavage site of TGFα before and after stimulation with PMA. Scale bars, 10 μm.

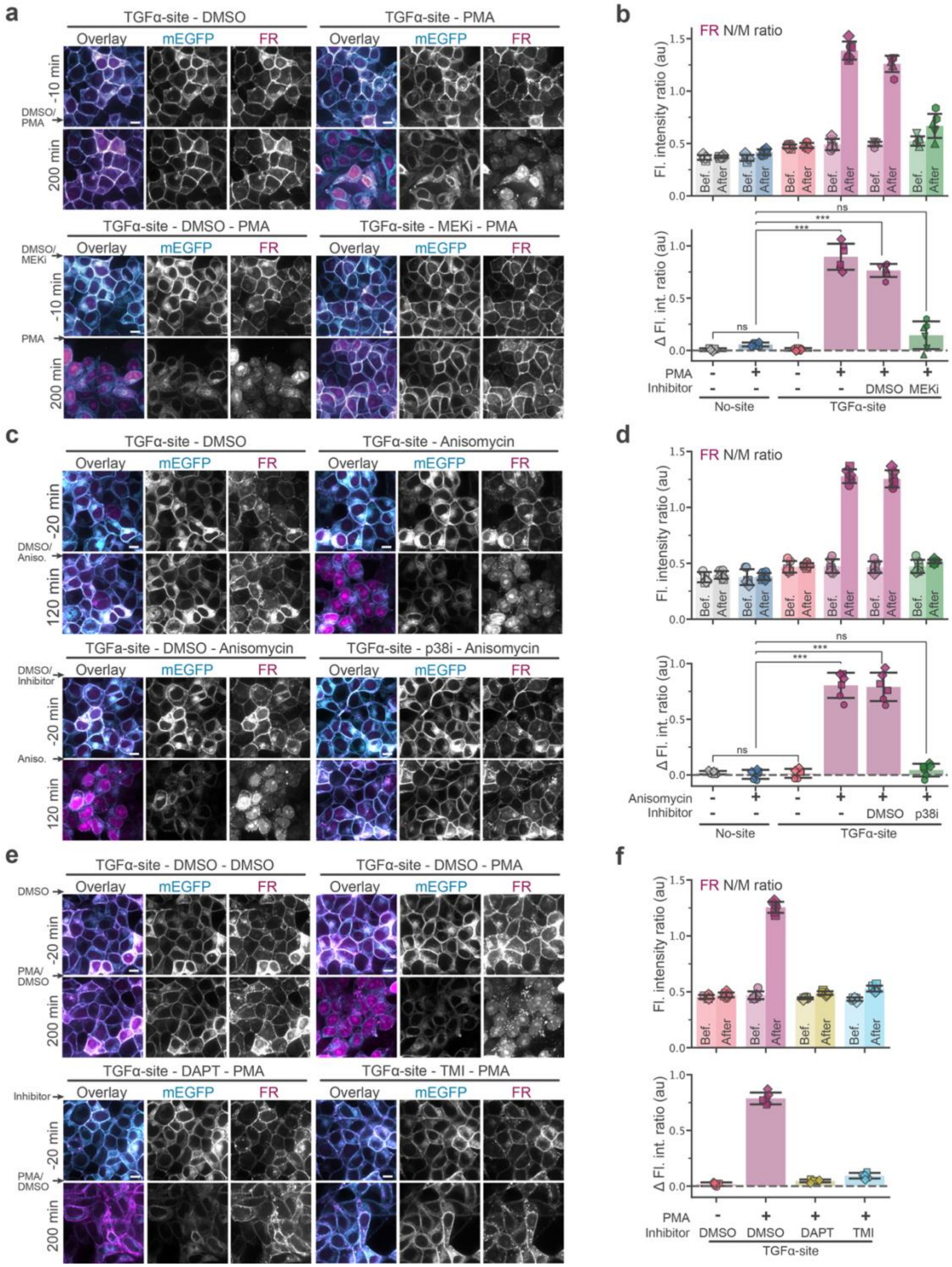


**Fig. S9: TGFα-site eNRGies specifically responds to ADAM17 activating and inhibiting stimuli. a, b,** Representative fluorescence images (**a**) and FR N/M ratio (top) and change in FR N/M ratio (bottom) (**b**) before (-10/-5min) and after (200-300 min) stimulation of cells expressing site-less (No-site) or TGFα-site eNRGies that were stimulated with DMSO or PMA at t = 0 and TGFα-site eNRGies expressing cells pretreated with DMSO or MEK inhibitor (MEKi) and stimulated with PMA at t = 0. **c, d,** Representative fluorescence images (**c**) and FR N/M ratio (top) and change in FR N/M ratio (bottom) (**d**) before (-20/-10 min) and after (100-200 min) stimulation of cells expressing site-less or TGFα-site eNRGies that were stimulated with DMSO or anisomycin at t = 0 and TGFα-site eNRGies expressing cells pretreated with DMSO or p38 inhibitor (p38i) and stimulated with anisomycin at t = 0. **e, f,** Representative fluorescence images (**e**) and FR N/M ratio (top) and change in FR N/M ratio (bottom) (**f**) before (-20/-10 min) and after (200 - 300 min) after stimulation of cells expressing TGFα-site eNRGies that were pretreated with DMSO, DAPT or TMI and stimulated with DMSO or PMA at t = 0. Bar and error bars represent the mean ± SD from 6 samples from three independent experiments identified by marker. Statistical analysis was conducted using a two-sided Welch's *t*-test. ns > 0.05, $*P \leq 0.05$, $**P \leq 0.01$ and $***P \leq 0.001$. Scale bars, 10 μm.

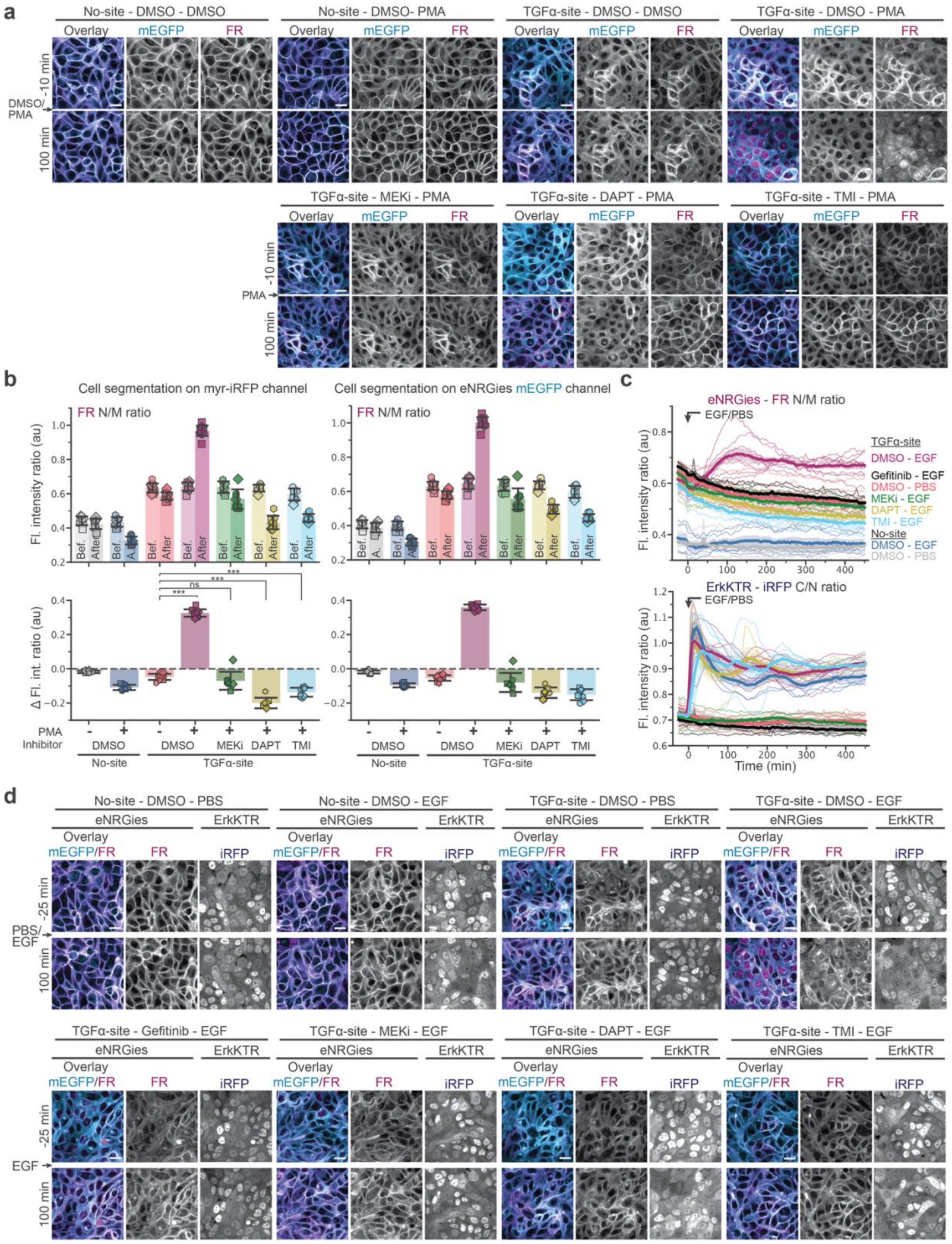

**Fig. S10: TGFα-site eNRGies specifically responds to ADAM17 activating and inhibiting stimuli in MCF10A cells. a,** Representative fluorescence images before and 100 min after stimulation of cells expressing site-less or TGFα-site eNRGies that were stimulated with DMSO or PMA at t = 0 and TGFα-site eNRGies expressing cells pretreated with DMSO, MEK inhibitor (MEKi), DAPT or PMI and stimulated with PMA at t = 0. **b**, FR N/M ratio (top) and change in FR N/M ratio (bottom) before (-10/-5min) and after (75-200 min) stimulation of cells expressing site-less or TGFα-site eNRGies that were stimulated with DMSO or PMA at t = 0 and TGFα-site eNRGies expressing cells pretreated with DMSO, MEK inhibitor (MEKi), DAPT or TMI and stimulated with PMA at t = 0. Cell body masks were generated using segmentation on the myr-iRFP channel (left) or the eNRGies mEGFP channel (right). Bar and error bars represent the mean ± SD from $N(\text{No-site}_{\text{DMSO}}) = 8$, $N(\text{No-site}_{\text{PMA}}) = 8$, $N(\text{TGF}\alpha\text{-site}_{\text{DMSO}}) = 12$, $N(\text{TGF}\alpha\text{-site}_{\text{PMA}}) = 12$, $N(\text{TGF}\alpha\text{-site}_{\text{MEKi-PMA}}) = 8$, $N(\text{TGF}\alpha\text{-site}_{\text{DAPT-PMA}}) = 8$, $N(\text{TGF}\alpha\text{-site}_{\text{TMI-PMA}}) = 8$ samples and four independent experiments identified by marker. Statistical analysis was conducted using a two-sided Welch's *t*-test. ns > 0.05, $*P \leq 0.05$, $**P \leq 0.01$ and $***P \leq 0.001$. **c, d,** eNRGies FR N/M and ErkKTR-iRFP C/N ratio time course (**c**) and fluorescence images (**d**) of MCF10A cells expressing site-less or TGFα-site eNRGies and the ErkKTR-iRFP that were stimulated with PBS or EGF at t = 0 and were pretreated with DMSO, Gefitinib, MEK inhibitor (MEKi), DAPT or TMI. Scale bars, 20 μm.

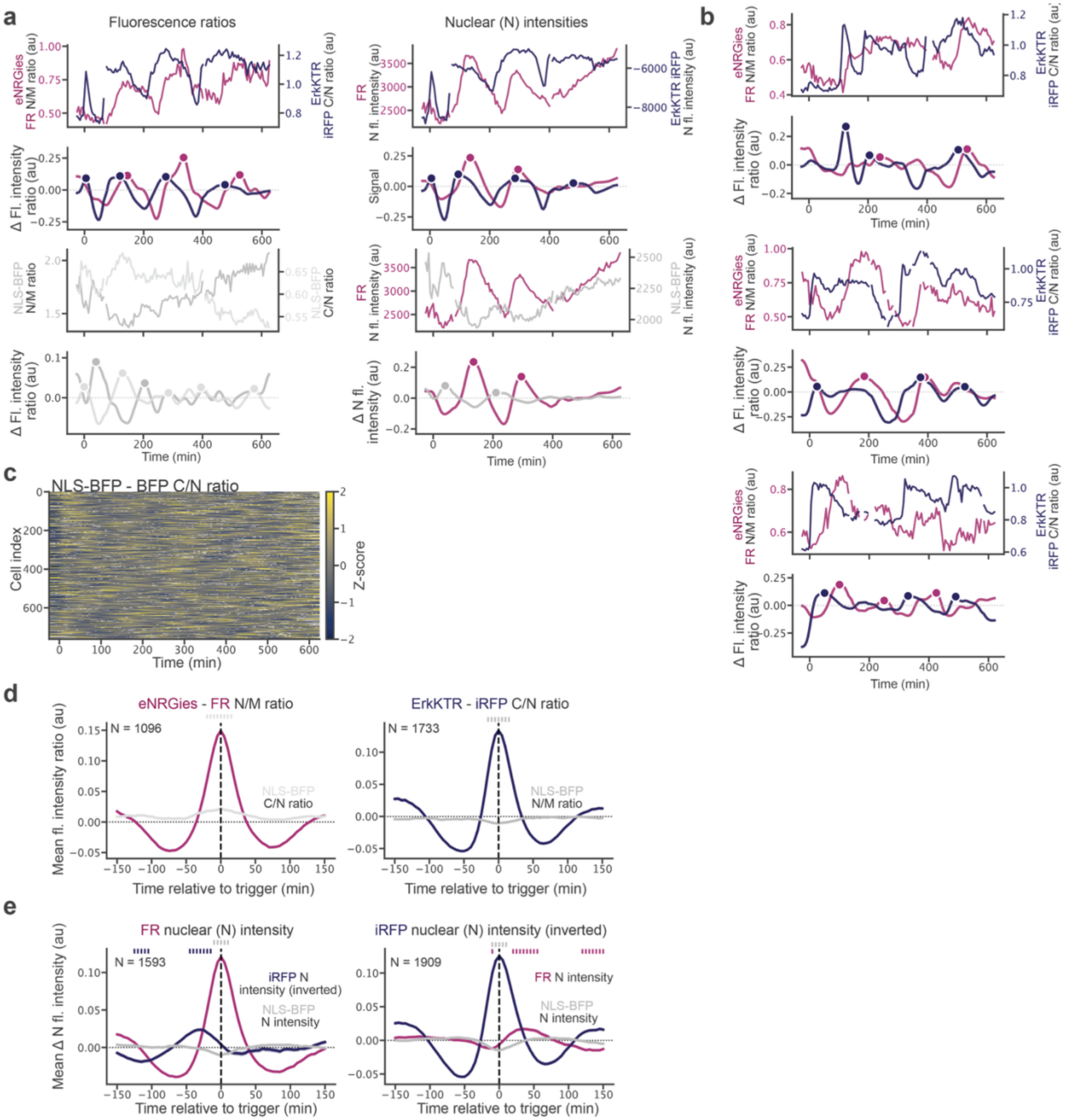


**Fig. S11: a,** Single-cell fluorescence intensity ratio (left) and nuclear fluorescence intensity traces (left) of an MCF10A cell expressing ADAM17 eNRGies and ErkKTR-iRFP showing activity pulses when exposed to EGF at t = 0 undergoing ADAM17 eNRGies and ErkKTR-iRFP pulses. **b,** Three additional representative single-cell fluorescence intensity ratio traces of an MCF10A cell expressing ADAM17 eNRGies and ErkKTR-iRFP showing activity pulses when exposed to EGF at t = 0**. a, b,** Top panel shows raw fluorescence intensity traces, lower panel shows smoothed and detrended traces with detected peaks marked (filled circles). **c,** Z-scored NLS-BFP C/N ratio per cell sorted according to the first peak in the eNRGies channel for 763 cells**.** White color indicates gaps in the tracking. **d,** Peak-triggered average of eNRGies FR N/M ratio (left) and ErkKTR-iRFP C/N ratio (right) peaks showing the aligned response of the control NLS-BFP C/N and N/M ratios. **e,** Peak-triggered average (baseline-substracted) of eNRGies FR nuclear intensity (left) and ErkKTR-iRFP inverted nuclear intensity (right) peaks showing the aligned response of the respective other sensor and the NLS-BFP control. **d, e,** Shading indicates SEM and N is the number of peak events averaged. Ticks indicate time points where the peak-triggered average of the co-recorded signal exceeds 70% of its peak amplitude.

### Supplementary Movie Legends:

**Movie S1: FR membrane fluorescence decreases for the Biosensor 1.0, but not the control protein.** Representative time series of cells expressing the initial biosensor design (mEGFP-TEVs-NRG-FR ) or a control protein (mEGFP-TEVs-CD28FL-FR) with the same extracellular and intracellular domains when exposed to PBS or 20U TEVp at time = 0. Snapshots are shown in Fig. 1c. Scale bars, 10 μm.

**Movie S2: Biosensors with additional NLS show nuclear localization of the ICD upon TEVp exposure, but a C-terminally located NLS leads to occasional mis-trafficking.** Representative time series of cells expressing the initial biosensor design (Biosensor 1.0) without additional NLS, or designs with an additional NLS at the C-terminal end (C-terminal NLS) or after the juxtamembrane domain (internal NLS) after exposure to 20 U TEVp at time = 0. Snapshots are shown in Fig. S2. Scale bars, 10 μm.

**Movie S3: Biosensors with long GS linkers or a helical linker are not efficiently cleaved by γ-secretase.** Representative time series of cells expressing a biosensor with additional internal NLS in which the original NRG linker sequence has been replaced with a GS linker or helical linker of varying length after exposure to 20 U TEVp at time = 0. Snapshots are shown in Fig. S3. Scale bars, 10 μm.

**Movie S4: Biosensors with the NRG transmembrane domain outperform those with an APP, Notch or CD28 transmembrane domain.** Representative time series of cells expressing the optimized biosensor with different transmembrane domains that are known γ-secretase substrates, NRG, Notch, APP or as control CD28 that is not a γ-secretase substrate after exposure to 20 U TEVp at time = 0. Snapshots are shown in Fig. S4. Scale bars, 10 μm.

**Movie S5: Biosensor response increases with increasing amounts of TEVp.** Representative time series of cells expressing TEVp or control eNRGies with no cleavage site after exposure to varying amounts of TEVp at time = 0. Snapshots are shown in Fig. 2a. Scale bars, 10 μm.

**Movie S6: eNRGies biosensors are dependent on secondary γ-secretase cleavage.** Representative time series of cells expressing the TEVp eNRGies after exposure to 20 U TEVp at time = 0 that were pretreated with or without γ-secretase inhibitors DAPT and Compound E (CompE). Snapshots are shown in Fig. 2e. Scale bars, 10 μm.

**Movie S7: Biosensor response recovers upon protease washout.** Representative time series of cells expressing the TEVp biosensor that were exposed to 20 U TEVp (top) or PBS (bottom) at time = 0 and were washed after 55 min of incubation to remove PBS or TEVp. Snapshots are shown in Fig. S5. Scale bars, 10 μm.

**Movie S8: eNRGies are a modular biosensor platform across multiple ECDs and ICDs.** Representative images of cells acquired in the ICD fluorescence channel expressing the TEVp eNRGies with different ECD and ICD domains after exposure to 20 U of TEVp at time = 0. Snapshots are shown in Fig. 3b. Scale bars, 10 μm.

**Movie S9**: **eNRGies can report on the activity of various soluble extracellular proteases.** Representative time series of cells expressing eNRGies biosensor with TEV-site, EK-site, FXa-site and MMP-9-site before and after exposure to 20 U TEVp, 32 U EK, 1 μg FXa or MMP-9 at time = 0. Snapshots are shown in Fig. 4b, e. Scale bars, 10 μm.

**Movie S10: TGFα-site eNRGies specifically responds to the ADAM17 stimuli PMA.** Representative fluorescence time series of cells expressing site-less or TGFα-site eNRGies that were stimulated with DMSO or PMA at time = 0 and TGFα-site eNRGies expressing cells pretreated with DMSO or MEK inhibitor (MEKi) and stimulated with PMA at t = 0. Snaphots are shown in Fig. S9. Scale bars, 10 μm.

**Movie S11: TGFα-site eNRGies specifically responds to the ADAM17 stimuli anisomycin.** Representative time series of cells expressing site-less or TGFα-site eNRGies that were stimulated with DMSO or anisomycin at t = 0 and TGFα-site eNRGies expressing cells pretreated with DMSO or p38 inhibitor (p38i) and stimulated with anisomycin at t = 0. Snapshots are shown in Fig. S9. Scale bars, 10 μm.

**Movie S12: TGFα-site eNRGies specifically responds to the ADAM17 inhibitor TMI.** Representative fluorescence time series of cells expressing TGFα-site eNRGies that were pretreated with DMSO, DAPT or TMI and stimulated with DMSO or PMA at t = 0. Snapshots are shown in Fig. S9. Scale bars, 10 µm.

**Movie S13: ADAM17 eNRGies reports on ADAM17 activity in HEK293T during mitosis**. Representative fluorescence time series of 6 different examples of each: HEK293T cells expressing a non-cleavable eNRGies or TGFα-site eNRGies and of ADAM17KO HEK293T expressing the TGFα-site eNRGies going through mitosis. Snapshots are shown in Fig. 6b, c. Scale bars, 10 µm.

**Movie S14: TGFα-site eNRGies specifically responds to ADAM17 activating and inhibiting stimuli in MCF10A cells.** Representative fluorescence time series of cells expressing site-less or TGFα-site eNRGies that were stimulated with DMSO or PMA at t = 0 and TGFα-site eNRGies expressing cells pretreated with DMSO, MEK inhibitor (MEKi), DAPT or PMI and stimulated with PMA at t = 0. Snapshots are shown in Fig. S10. Scale bars, 20 µm.

**Movie S15: TGFα-site eNRGies specifically responds to EGF stimulation in MCF10A cells.** Fluorescence time series of MCF10A cells expressing site-less or TGFα-site eNRGies and the ErkKTR-iRFP that were stimulated with PBS or EGF at t = 0 and were pretreated with DMSO, Gefitinib, MEK inhibitor (MEKi), DAPT or TMI. Snapshots are shown in Fig. S10. Scale bars, 20 µm.

**Movie S16: eNRGies biosensor reveals dynamics of ADAM17 activity in response to EGF stimulation.** Representative fluorescence time series of 5 tracked MCF10A cell expressing ADAM17 eNRGies and ErkKTR-iRFP showing activity pulses when exposed to EGF at t = 0. Tracked cell outline is shown in yellow. Snapshots of cell 3 are shown in Fig. 7f. Scale bars, 20 µm.

### Supplementary Tables

**Supplementary Table S1.** Plasmids used in this study with key plasmids highlighted in bold. With the exception of pQC110 all plasmids were constructed in this study.

| Name | # Addgene ID | Short name | Description of expressed proteins | Cleavage site[1] | Linker sequence | Relevant figures/ movies |
|---|---|---|---|---|---|---|
| **Nuclear and membrane marker, ErkKTR** | | | | | | |
| **pBR181** | 259190 | myr-iRFP NLS-BFP | myr-iRFP- P2A-NLS-TagBFP | - | - | 1-7 |
| **pBR227** | 259191 | myr-iRFP NLS-FusionRed | myr-iRFP- P2A-NLS-FusionRed | - | - | 3, S6, Movie S8 |
| **pBR254** | 259192 | myr-FusionRed NLS-TagBFP | myr-FusionRed-P2A- NLS-TagBFP | - | - | 3, S6, Movie S8 |
| pQC110 | | ErkKTR-iRFP NLS-BFP | ErkKTR-iRFP-P2A-NLS-TagBFP | | | 7d-i, S10, S11, Movie S15, S16 |
| **Biosensor plasmids** | | | | | | |
| pBR20 | | | CD4-mEGFP-TEVs-CD28-FusionRed | LGKS-ENLYFQ/M-GGSG (TEV-site) | | 1c-e, Movie S1 |
| pBR100 | | | CD4-mEGFP-TEVs-NRGlinker-NRG-FusionRed | LGKS-ENLYFQ/M-GGSG (TEV-site) | MEAEELYQKR (from NRG) | 1c-e, g, S2, Movie S1, S2 |
| pBR126 | | | CD4-mEGFP-TEVs-NRGlinker-NRG-FusionRed-NLS | LGKS-ENLYFQ/M-GGSG (TEV-site) | MEAEELYQKR (from NRG) | 1g, S2, Movie S2 |
| pBR127 | | | CD4-mEGFP-TEVs-NRGlinker-NRG-NLS-FusionRed | LGKS-ENLYFQ/M-GGSG (TEV-site) | MEAEELYQKR (from NRG) | 1g, S2, Movie S2 |
| pBR166 | | | CD4-mEGFP-TEVs-10AAGS-NRG-NLS-FusionRed | LGKS-ENLYFQ/M (TEV-site) | 2x(GGGGS) | 1h, S3, Movie S3 |
| **pBR167** | 259193 | TEV-site | CD4-mEGFP-TEVs-20AAGS-NRG-NLS-FusionRed | LGKS-ENLYFQ/M (TEV-site) | 4x(GGGGS) | 1h, 2-4, S3-S9, Movie S3-S9 |
| pBR242 | | | CD4-mEGFP-TEVs-30AAGS-NRG-NLS-FusionRed | LGKS-ENLYFQ/M (TEV-site) | 5x(GS)+4x(GGGGS) | 1h, S3, Movie S3 |
| pBR243 | | | CD4-mEGFP-TEVs-40AAGS-NRG-NLS-FusionRed | LGKS-ENLYFQ/M (TEV-site) | 10x(GS)+4x(GGGGS) | 1h, S3, Movie S3 |

| | | | | | | |
|---|---|---|---|---|---|---|
| pBR244 | | | CD4-mEGFP-TEVs-50AAGS-NRG-NLS-FusionRed | LGKS-ENLYFQ/M (TEV-site) | 15x(GS)+4x(GGGGS) | 1h, S3, Movie S3 |
| pBR174 | | | CD4-mEGFP-TEVs-30AAhelicallinker-NRG-NLS-FusionRed | LGKS-ENLYFQ/M (TEV-site) | 6x(EAAAK) | S3, Movie S3 |
| pBR175 | | | CD4-mEGFP-TEVs-20AAhelicallinker-NRG-NLS-FusionRed | LGKS-ENLYFQ/M (TEV-site) | 4x(EAAAK) | S3, Movie S3 |
| pBR176 | | | CD4-mEGFP-TEVs-10AAhelicallinker-NRG-NLS-FusionRed | LGKS-ENLYFQ/M (TEV-site) | 2x(EAAAK) | S3, Movie S3 |
| pBR194 | | | CD4-mEGFP-TEVs-20AAGS-NotchTM-NLS-FusionRed | LGKS-ENLYFQ/M (TEV-site) | 4x(GGGGS) | 1i, S4, Movie S4 |
| pBR195 | | | CD4-mEGFP-TEVs-20AAGS-APPTM-NLS-FusionRed | LGKS-ENLYFQ/M (TEV-site) | 4x(GGGGS) | 1i, S4, Movie S4 |
| pBR200 | | | CD4-mEGFP-TEVs-20AAGS-CD28TM-NLS-FusionRed | LGKS-ENLYFQ/M (TEV-site) | 4x(GGGGS) | 1i, S4, Movie S4 |
| **pBR222** | 259194 | | CD4-SEAP-2xmyc-TEVs-20AAGS-NRG-NLS-FusionRed | LGKS-ENLYFQ/M (TEV-site) | 4x(GGGGS) | 3, S6, Movie S8 |
| **pBR224** | 259195 | | CD4-SEAP-2xmyc-TEVs-20AAGS-NRG-NLS-Clover | LGKS-ENLYFQ/M (TEV-site) | 4x(GGGGS) | 3, S6, Movie S8 |
| **pBR225** | 259196 | | CD4-SEAP-2xmyc-TEVs-20AAGS-NRG-NLS-mTagBFP | LGKS-ENLYFQ/M (TEV-site) | 4x(GGGGS) | 3, S6, Movie S8 |
| **pBR252** | 259197 | | CD4-SEAP-2xmyc-TEVs-20AAGS-NRG-NLS-Halotag | LGKS-ENLYFQ/M (TEV-site) | 4x(GGGGS) | 3, S6, Movie S8 |
| **pBR253** | 259198 | | CD4-SEAP-2xmyc-TEVs-20AAGS-NRG-NLS-iRFP | LGKS-ENLYFQ/M (TEV-site) | 4x(GGGGS) | 3, S6, Movie S8 |
| **pBR188** | 259199 | EK-site | CD4-mEGFP-EKs-20AAGS-NRG-NLS-FusionRed | LGKSGS-DDDDK/M (Ek-site) | 4x(GGGGS) | 4b-d, Movie S9 |

| | | | | | | |
|---|---|---|---|---|---|---|
| **pBR189** | 259200 | Fxa-site | CD4-mEGFP-FXas-20AAGS-NRG-NLS-FusionRed | LGKSGS-IDGR/M (Fxa-site) | 4x(GGGGS) | 4b-d, Movie S9 |
| **pBR198** | 259201 | MMP-9-site | CD4-mEGFP-MMP9s-20AAGS-NRG-NLS-FusionRed | LGKSGS-KGPRS/LSGK (MMP-9-site) | 4x(GGGGS) | 4e-g, Movie S9 |
| pBR186 | | ADAMtide | CD4-mEGFP-ADAMtide-16AAGS-NRG-NLS-FusionRed | LGKS-PRAEA/LKGG | G-3x(GGGGS) | S7 |
| pBR187 | | TENtide | CD4-mEGFP-TENtide-16AAGS-NRG-NLS-FusionRed | LGKS-PRYEA/YKMG | G-3x(GGGGS) | S7 |
| pBR196 | | TNFtide | CD4-mEGFP-TNFtide-16AAGS-NRG-NLS-FusionRed | LGKS-PLAQA/VRSS | G-3x(GGGGS) | S7 |
| pBR197 | | TACEtide | CD4-mEGFP-TACEtide-16AAGS-NRG-NLS-FusionRed | LGKS-PRAAA/VKSP | G-3x(GGGGS) | S7 |
| **pBR231** | 259202 | No-site/ Control eNRGies | CD4-mEGFP-20AAGS-NRG-NLS-FusionRed | - | 4x(GGGGS) | 5-7, S8-S11, Movie S10-S15 |
| pBR234 | | EGF-site | CD4-mEGFP-EGFs-10AAGS-NRG-NLS-FusionRed | LGKS-WWELR/HAGH (#P01133, AA1019-1027 of human EGF) | 2x(GGGGS) | 5b, c, S8 |
| **pBR235** | 259203 | TGFa-site/ ADAM17 eNRGies | CD4-mEGFP-TGFas-10AAGS-NRG-NLS-FusionRed | ADLLA/VVAA (#P01135, AA85-93 of human TGFa) | 2x(GGGGS) | 5-7, S8-S11, Movie S10-S16 |
| pBR237 | | AREG-site | CD4-mEGFP-AREGas-10AAGS-NRG-NLS-FusionRed | RCGEKSM/KT (#P15514, AA181-188 of human AREG) | 2x(GGGGS) | 5b, c, S8 |

[1] The predicted cleavage site is indicated with "/".

**Supplementary Table S2.** List of protein sequences used in this study.

| Domain names | Description | Amino acid sequence |
|---|---|---|
| myr | myristoylation sequence - DPVAT linker | MGSSKSKPKDASQRRR-DPVAT |
| P2A | P2A sequence - AT linker | GSGATNFSLLKQAGDVEENPGP-AT |
| NLS for nuclear marker | linker-SV40 NLS - linker | MA-PKKKRKV-RYPAFLYKVAT |

| | | |
|---|---|---|
| ErkKTR | ErkKTR - linker | MKGRKPRDLELPLSPSLLGGQGPERTPGSGTSSGLQAPGPALSP SKRSGLEDPATPSKKPRTPSVSSRLERLTLQSSFQFPSTS-TRQVEQGRWTVDPVAT |
| CD4 | mouse CD4 signal sequence - GGGGS linker | MCRAISLRRLLLLLLQLSQLLAVTQG-GGGGS |
| CD28 | human CD28 (#P10747, AA19-220) -GGG linker | NKILVKQSPMLVAYDNAVNLSCKYSYNLFSREFRASLHKGLDS AVEVCVVYGNYSQQLQVYSKTGFNCDGKLGNESVTFYLQNLY VNQTDIYFCKIEVMYPPPYLDNEKSNGTIIHVKGKHLCPSPLFPG PSKPFWVLVVVGGVLACYSLLVTVAFIIFWVRSKRSRLLHSDY MNMTPRRPGPTRKHYQPYAPPRDFAAYRS-GGG |
| NRG | NRG TMD and juxtamembrane domain (Rat NRG III 1a) - GGG linker | VLTITGICIALLVVGIMCVVAYC-KTKKQRQK-GGG |
| NLS in eNRgies | SV40 NLS - GS linker | PKKKRKV-GS |
| Notch | human Notch 1 TMD (#P46531,AA1736-1756)-NRG juxtamembrane domain - GGG linker | FMYVAAAAFVLLFFVGCGVLLS-KTKKQRQK-GGG |
| APP | human APP TMD (#P05067, AA702-723)-NRG juxtamembrane domain - GGG linker | IIGLMVGGVVIATVIVITLVML-KTKKQRQK-GGG |
| CD28TM | human CD28 TMD (#P10747,AA153-179)-NRG juxtamembrane domain - GGG linker | FWVLVVVGGVLACYSLLVTVAFIIFWV-KTKKQRQK-GGG |
| SEAP | secreted alkaline phosphatase, human (#P05187,AA6-511) | MLLLLLLLGLRLQLSLGIIPVEEENPDFWNREAAEALGAAKKLQ PAQTAAKNLIIFLGDGMGVSTVTAARILKGQKKDKLGPEIPLAM DRFPYVALSKTYNVDKHVPDSGATATAYLCGVKGNFQTIGLSA AARFNQCNTTRGNEVISVMNRAKKAGKSVGVVTTTRVQHASP AGTYAHTVNRNWYSDADVPASARQEGCQDIATQLISNMDIDVI LGGGRKYMFRMGTPDPEYPDDYSQGGTRLDGKNLVQEWLAK RQGARYVWNRTELMQASLDPSVTHLMGLFEPGDMKYEIHRDS TLDPSLMEMTEAALRLLSRNPRGFFLFVEGGRIDHGHHESRAYR ALTETIMFDDAIERAGQLTSEEDTLSLVTADHSHVFSFGGYPLR GSSIFGLAPGKARDRKAYTVLLYGNGPGYVLKDGARPDVTESE SGSPEYRQQSAVPLDEETHAGEDVAVFARGPQAHLVHGVQEQ TFIAHVMAFAACLEPYTACDLAPPAGTTDAAHPG |
| 2xmyc | Linker - 2x myc tag | GGSG-EQKLISEEDLEQKLISEEDL |
| FusionRed (FR) | FusionRed | MVSELIKENMPMKLYMEGTVNNHHFKCTSEGEGKPYEGTQTM RIKVVEGGPLPFAFDILATSFMYGSRTFIKHPPGIPDFFKQSFPEG FTWERVTTYEDGGVLTATQDTSLQDGCLIYNVKVRGVNFPAN GPVMQKKTLGWEASTETMYPADGGLEGACDMALKLVGGGHL ICNLETTYRSKKPATNLKMPGVYNVDHRLERIKEADDETYVEQ HEVAVARYSTGGAGDGGKAATR |
| mEGFP | mEGFP | MVSKGEELFTGVVPILVELDGDVNGHKFSVSGEGEGDATYGKL TLKFICTTGKLPVPWPTLVTTLTYGVQCFSRYPDHMKQHDFFKS AMPEGYVQERTIFFKDDGNYKTRAEVKFEGDTLVNRIELKGIDF KEDGNILGHKLEYNYNSHNVYIMADKQKNGIKVNFKIRHNIED GSVQLADHYQQNTPIGDGPVLLPDNHYLSTQSKLSKDPNEKRD HMVLLEFVTAAGITLGMDELYK |

| | | |
|---|---|---|
| Clover | Clover | MVSKGEELFTGVVPILVELDGDVNGHKFSVRGEGEGDATNGKL TLKFICTTGKLPVPWPTLVTTFGYGVACFSRYPDHMKQHDFFK SAMPEGYVQERTISFKDDGTYKTRAEVKFEGDTLVNRIELKGID FKEDGNILGHKLEYNFNSHNVYITADKQKNGIKANFKIRHNVE DGSVQLADHYQQNTPIGDGPVLLPDNHYLSHQSALSKDPNEKR DHMVLLEFVTAA |
| TagBFP | TagBFP | MSELIKENMHMKLYMEGTVDNHHFKCTSEGEGKPYEGTQTMR IKVVEGGPLPFAFDILATSFLYGSKTFINHTQGIPDFFKQSFPEGF TWERVTTYEDGGVLTATQDTSLQDGCLIYNVKIRGVNFTSNGP VMQKKTLGWEAFTETLYPADGGLEGRNDMALKLVGGSHLIAN AKTTYRSKKPAKNLKMPGVYYVDYRLERIKEANNETYVEQHE VAVARYCDLPSKLGHKLN |
| Halotag | HaloTag | EIGTGFPFDPHYVEVLGERMHYVDVGPRDGTPVLFLHGNPTSS YVWRNIIPHVAPTHRCIAPDLIGMGKSDKPDLGYFFDDHVRFM DAFIEALGLEEVVLVIHDWGSALGFHWAKRNPERVKGIAFMEFI RPIPTWDEWPEFARETFQAFRTTDVGRKLIIDQNVFIEGTLPMG VVRPLTEVEMDHYREPFLNPVDREPLWRFPNELPIAGEPANIVA LVEEYMDWLHQSPVPKLLFWGTPGVLIPPAEAARLAKSLPNCK AVDIGPGLNLLQEDNPDLIGSEIARWLSTLEISG |
| iRFP | iRFP | MAEGSVARQPDLLTCDDEPIHIPGAIQPHGLLLALAADMTIVAG SDNLPELTGLAIGALIGRSAADVFDSETHNRLTIALAEPGAAVG APITVGFTMRKDAGFIGSWHRHDQLIFLELEPPQRDVAEPQAFF RRTNSAIRRLQAAETLESACAAAAQEVRKITGFDRVMIYRFASD FSGEVIAEDRCAEVESKLGLHYPASTVPAQARRLYTINPVRIIPDI NYRPVPVTPDLNPVTGRPIDLSFAILRSVSPVHLEFMRNIGMHGT MSISILRGERLWGLIVCHHRTPYYVDLDGRQACELVAQVLAWQ IGVMEE |

**Supplementary Table S3.** Cell lines used in this study.

| Cell line | Transgenes | Relevant figures/ movies | Source |
|---|---|---|---|
| **Standard HEK293T** | | | |
| HEK293T | Parental cell line | None | Toettcher Lab |
| myr-iRFP_NLS-BFP | pBR181 (monoclonal) | None | This study |
| myr-iRFP_NLS-FusionRed | pBR227 (monoclonal) | None | This study |
| myr-FusionRed_NLS-TagBFP | pBR254 (monoclonal) | None | This study |
| CD4-mEGFP-TEVs-CD28-FusionRed, myr-iRFP NLS-BFP | pBR20 (polyclonal), pBR181 (monoclonal) | 1c-e, Movie S1 | This study |
| CD4-mEGFP-TEVs-NRGlinker-NRG-FusionRed, myr-iRFP NLS-BFP | pBR100 (polyclonal), pBR181 (monoclonal) | 1c-e, g, S2, Movie S1, S2 | This study |
| CD4-mEGFP-TEVs-NRGlinker-NRG-FusionRed-NLS, myr-iRFP NLS-BFP | pBR126 (polyclonal), pBR181 (monoclonal) | 1g, S2, Movie S2 | This study |
| CD4-mEGFP-TEVs-NRGlinker-NRG-NLS-FusionRed, myr-iRFP NLS-BFP | pBR127 (polyclonal), pBR181 (monoclonal) | 1g, S2, Movie S2 | This study |
| CD4-mEGFP-TEVs-10AAGS-NRG-NLS-FusionRed, myr-iRFP NLS-BFP | pBR166 (polyclonal), pBR181 (monoclonal) | 1h, S3, Movie S3 | This study |
| CD4-mEGFP-TEVs-20AAGS-NRG-NLS-FusionRed, myr-iRFP NLS-BFP | pBR167 (polyclonal), pBR181 (monoclonal) | 1h, S3-S7, 2-4, Movie S3-9 | This study |
| CD4-mEGFP-TEVs-20AAGS-NRG-NLS-FusionRed monoclonal, myr-iRFP NLS-BFP | pBR167 (monoclonal), pBR181 (monoclonal) | 2g | This study |

| | | | |
|---|---|---|---|
| CD4-mEGFP-TEVs-30AAGS-NRG-NLS-FusionRed, myr-iRFP NLS-BFP | pBR242 (polyclonal), pBR181 (monoclonal) | 1h, S3, Movie S3 | This study |
| CD4-mEGFP-TEVs-40AAGS-NRG-NLS-FusionRed, myr-iRFP NLS-BFP | pBR243 (polyclonal), pBR181 (monoclonal) | 1h, S3, Movie S3 | This study |
| CD4-mEGFP-TEVs-50AAGS-NRG-NLS-FusionRed, myr-iRFP NLS-BFP | pBR244 (polyclonal), pBR181 (monoclonal) | 1h, S3, Movie S3 | This study |
| CD4-mEGFP-TEVs-30AAhelicallinker-NRG-NLS-FusionRed, myr-iRFP NLS-BFP | pBR174 (polyclonal), pBR181 (monoclonal) | S3, Movie S3 | This study |
| CD4-mEGFP-TEVs-20AAhelicallinker-NRG-NLS-FusionRed, myr-iRFP NLS-BFP | pBR175 (polyclonal), pBR181 (monoclonal) | S3, Movie S3 | This study |
| CD4-mEGFP-TEVs-10AAhelicallinker-NRG-NLS-FusionRed, myr-iRFP NLS-BFP | pBR176 (polyclonal), pBR181 (monoclonal) | S3, Movie S3 | This study |
| CD4-mEGFP-TEVs-20AAGS-NotchTM-NLS-FusionRed, myr-iRFP NLS-BFP | pBR194 (polyclonal), pBR181 (monoclonal) | 1i, S4, Movie S4 | This study |
| CD4-mEGFP-TEVs-20AAGS-APPTM-NLS-FusionRed, myr-iRFP NLS-BFP | pBR195 (polyclonal), pBR181 (monoclonal) | 1i, S4, Movie S4 | This study |
| CD4-mEGFP-TEVs-20AAGS-CD28TM-NLS-FusionRed, myr-iRFP NLS-BFP | pBR200 (polyclonal), pBR181 (monoclonal) | 1i, S4, Movie S4 | This study |
| CD4-SEAP-2xmyc-TEVs-20AAGS-NRG-NLS-FusionRed, myr-iRFP NLS-BFP | pBR222 (polyclonal), pBR181 (monoclonal) | 3, S6, Movie S8 | This study |
| CD4-SEAP-2xmyc-TEVs-20AAGS-NRG-NLS-Clover, myr-iRFP NLS-BFP | pBR224 (polyclonal), pBR181 (monoclonal) | 3, S6, Movie S8 | This study |
| CD4-SEAP-2xmyc-TEVs-20AAGS-NRG-NLS-mTagBFP, myr-iRFP NLS-FusionRed | pBR225 (polyclonal), pBR227 (monoclonal) | 3, S6, Movie S8 | This study |
| CD4-SEAP-2xmyc-TEVs-20AAGS-NRG-NLS-Halotag, myr-iRFP NLS-BFP | pBR252 (polyclonal), pBR181 (monoclonal) | 3, S6, Movie S8 | This study |
| CD4-SEAP-2xmyc-TEVs-20AAGS-NRG-NLS-iRFP, myr-iRFP NLS-BFP | pBR253 (polyclonal), pBR254 (monoclonal) | 3, S6, Movie S8 | This study |
| Ek-site, myr-iRFP NLS-BFP | pBR188 (polyclonal), pBR181 (monoclonal) | 4b-d, Movie S9 | This study |
| Fxa-site, myr-iRFP NLS-BFP | pBR189 (polyclonal), pBR181 (monoclonal) | 4b-d, Movie S9 | This study |
| MMP-9-site, myr-iRFP NLS-BFP | pBR198 (polyclonal), pBR181 (monoclonal) | 4e-g, Movie S9 | This study |
| No-site, myr-iRFP NLS-BFP | pBR231 (polyclonal), pBR181 (monoclonal) | 5, S8-S9, Movie S10-12 | This study |
| No-site (monoclonal), myr-iRFP NLS-BFP | pBR231 (monoclonal), pBR181 (monoclonal) | 5, Movie S13 | This study |
| EGF-site, myr-iRFP NLS-BFP | pBR234 (polyclonal), pBR181 (monoclonal) | 5b, c, S8 | This study |
| TGFa-site, myr-iRFP NLS-BFP | pBR235 (polyclonal), pBR181 (monoclonal) | 5, S8-S9, Movie S10-12 | This study |
| TGFa-site (monoclonal), myr-iRFP NLS-BFP | pBR235 (monoclonal), pBR181 (monoclonal) | 5, Movie S13 | This study |
| AREG-site, myr-iRFP NLS-BFP | pBR237 (polyclonal), pBR181 (monoclonal) | 5b, c, S8 | This study |
| **HEK293T KO cell lines** | | | |
| HEK293T WT | Parental cell line | None | Veit Hornung (LMU) |
| HEK293T ADAM10 KO | Parental cell line | None | Veit Hornung (LMU) |

| | | | |
|---|---|---|---|
| HEK293T ADAM17 KO | Parental cell line | None | Veit Hornung (LMU) |
| HEK293T ADAM1/ADAM17 DKO | Parental cell line | None | Veit Hornung (LMU) |
| HEK293T WT, myr-iRFP NLS-BFP | pBR181 (monoclonal) | None | This study |
| HEK293T ADAM10 KO, myr-iRFP NLS-BFP | pBR181 (monoclonal) | None | This study |
| HEK293T ADAM17 KO, myr-iRFP NLS-BFP | pBR181 (monoclonal) | None | This study |
| HEK293T ADAM1/ADAM17 DKO, myr-iRFP NLS-BFP | pBR181 (monoclonal) | None | This study |
| HEK293T WT, myr-iRFP NLS-BFP, ADAMtide | pBR186 (polyclonal), pBR181 (monoclonal) | S7 | This study |
| HEK293T ADAM10 KO, myr-iRFP NLS-BFP, ADAMtide | pBR186 (polyclonal), pBR181 (monoclonal) | S7 | This study |
| HEK293T ADAM17 KO, myr-iRFP NLS-BFP, ADAMtide | pBR186 (polyclonal), pBR181 (monoclonal) | S7 | This study |
| HEK293T ADAM1/ADAM17 DKO, myr-iRFP NLS-BFP, ADAMtide | pBR186 (polyclonal), pBR181 (monoclonal) | S7 | This study |
| HEK293T WT, myr-iRFP NLS-BFP, TENtide | pBR187 (polyclonal), pBR181 (monoclonal) | S7 | This study |
| HEK293T ADAM10 KO, myr-iRFP NLS-BFP, TENtide | pBR187 (polyclonal), pBR181 (monoclonal) | S7 | This study |
| HEK293T ADAM17 KO, myr-iRFP NLS-BFP, TENtide | pBR187 (polyclonal), pBR181 (monoclonal) | S7 | This study |
| HEK293T ADAM1/ADAM17 DKO, myr-iRFP NLS-BFP, TENtide | pBR187 (polyclonal), pBR181 (monoclonal) | S7 | This study |
| HEK293T WT, myr-iRFP NLS-BFP, TNFtide | pBR196 (polyclonal), pBR181 (monoclonal) | S7 | This study |
| HEK293T ADAM10 KO, myr-iRFP NLS-BFP, TNFtide | pBR196 (polyclonal), pBR181 (monoclonal) | S7 | This study |
| HEK293T ADAM17 KO, myr-iRFP NLS-BFP, TNFtide | pBR196 (polyclonal), pBR181 (monoclonal) | S7 | This study |
| HEK293T ADAM1/ADAM17 DKO, myr-iRFP NLS-BFP, TNFtide | pBR196 (polyclonal), pBR181 (monoclonal) | S7 | This study |
| HEK293T WT, myr-iRFP NLS-BFP, TACEtide | pBR197 (polyclonal), pBR181 (monoclonal) | S7 | This study |
| HEK293T ADAM10 KO, myr-iRFP NLS-BFP, TACEtide | pBR197 (polyclonal), pBR181 (monoclonal) | S7 | This study |
| HEK293T ADAM17 KO, myr-iRFP NLS-BFP, TACEtide | pBR197 (polyclonal), pBR181 (monoclonal) | S7 | This study |
| HEK293T ADAM1/ADAM17 DKO, myr-iRFP NLS-BFP, TACEtide | pBR197 (polyclonal), pBR181 (monoclonal) | S7 | This study |
| HEK293T WT, myr-iRFP NLS-BFP, No site | pBR231 (polyclonal), pBR181 (monoclonal) | 5d, S8 | This study |
| HEK293T ADAM10 KO, myr-iRFP NLS-BFP, No site | pBR231 (polyclonal), pBR181 (monoclonal) | 5d, S8 | This study |
| HEK293T ADAM17 KO, myr-iRFP NLS-BFP, No site | pBR231 (polyclonal), pBR181 (monoclonal) | 5d, S8 | This study |
| HEK293T ADAM1/ADAM17 DKO, myr-iRFP NLS-BFP, No site | pBR231 (polyclonal), pBR181 (monoclonal) | 5d, S8 | This study |
| HEK293T WT, myr-iRFP NLS-BFP, TGFa-site | pBR235 (polyclonal), pBR181 (monoclonal) | 5d, S8 | This study |

| | | | |
|---|---|---|---|
| HEK293T ADAM10 KO, myr-iRFP NLS-BFP, TGFa-site | pBR235 (polyclonal), pBR181 (monoclonal) | 5d, S8 | This study |
| HEK293T ADAM17 KO, myr-iRFP NLS-BFP, TGFa-site | pBR235 (polyclonal), pBR181 (monoclonal) | 5d, S8 | This study |
| HEK293T ADAM1/ADAM17 DKO, myr-iRFP NLS-BFP, TGFa-site | pBR235 (polyclonal), pBR181 (monoclonal) | 5d, S8 | This study |
| **MCF10A cell lines** | | | |
| MCF10A 5E | Parental cell line | None | Toettcher Lab |
| myr-iRFP NLS-BFP | pBR181 (monoclonal) | None | This study |
| ErkKTR-iRFP NLS-BFP | pQC110 (monoclonal) | None | This study |
| No-site, myr-iRFP NLS-BFP | pBR231 (monoclonal, pBR181 (monoclonal) | 7b,c, S10, Movie S14 | This study |
| TGFa-site, myr-iRFP NLS-BFP | pBR235 (monoclonal, pBR181 (monoclonal) | 7b,c, S10, Movie S14 | This study |
| No-site, ErkKTR-iRFP NLS-BFP | pBR231 (monoclonal, pQC110 (monoclonal) | 7d,e, S10, Movie S15 | This study |
| TGFa-site,ErkKTR-iRFP NLS-BFP | pBR235 (monoclonal, pQC110 (monoclonal) | 7d-i, S10, S11, Movie S15, S16 | This study |
